# Reusable rule-based cell cycle model explains compartment-resolved dynamics of 16 observables in RPE-1 cells

**DOI:** 10.1101/2023.05.04.539349

**Authors:** Paul F. Lang, David R. Penas, Julio R. Banga, Daniel Weindl, Bela Novak

## Abstract

The mammalian cell cycle is regulated by a well-studied but complex biochemical reaction system. Computational models provide a particularly systematic and systemic description of the mechanisms governing mammalian cell cycle control. By combining both state-of-the-art multiplexed experimental methods and powerful computational tools, this work aims at improving on these models along four dimensions: model structure, validation data, validation methodology and model reusability.

We developed a comprehensive model structure of the full cell cycle that qualitatively explains the behaviour of human retinal pigment epithelial-1 cells. To estimate the model parameters, time courses of eight cell cycle regulators in two compartments were reconstructed from single cell snapshot measurements. After optimisation with a parallel global optimisation metaheuristic we obtained excellent agreements between simulations and measurements. The PEtab specification of the optimisation problem facilitates reuse of model, data and/or optimisation results.

Future perturbation experiments will improve parameter identifiability and allow for testing model predictive power. Such a predictive model may aid in drug discovery for cell cycle-related disorders.

**Author Summary:** While there are numerous cell cycle models in the literature, mammalian cell cycle models typically suffer from four limitations. Firstly, the descriptions of biological mechanisms are often overly complicated yet insufficiently comprehensive and detailed. Secondly, there is a lack of experimental data to validate the model. Thirdly, inadequate parameter estimation procedures are used. Lastly, there is no standardized description of the model and/or optimization problem.

To overcome these limitations, we combine best-in-class technology to address all four simultaneously. We use a rule-based model description to provide a concise and less error-prone representation of complex biology. By applying trajectory reconstruction algorithms to existing data from highly multiplexed immunofluorescence measurements, we obtained a rich dataset for model validation. Using a parallel global metaheuristic for parameter estimation allowed us to bring simulations and data in very good agreement. To maximize reproducibility and reusability of our work, the results are available in three popular formats: BioNetGen, SBML, and PEtab.

Our model is generalizable to many healthy and transformed cell types. The PEtab specification of the optimization problem makes it straightforward to re-optimize the parameters for other cell lines. This may guide hypotheses on cell type-specific regulation of the cell cycle, potentially with clinical relevance.

## Introduction

Scientific research can be seen as a collaborative process of continuously refining models of the world. Often, these refinements are driven by acquisition of new types of data, more accurate data, better data analysis methods or novel ways to describe, communicate, explain or interpret data and knowledge. In case of systems biology, we now have access to highly multiplexed measurements of biopolymer composition at single cell resolution [1–3], methods to analyse [4–11], and formats to communicate and interpret these data [12–15]. However, these methods and tools have not yet been exploited to refine existing models of the mammalian cell cycle control system. Through an intricate interplay of protein synthesis, degradation, phosphorylation and complexation, this cell cycle control system (1) ensures strictly alternating replication and segregation of DNA, (2) coordinates these processes with doubling of all other cellular components, (3) checks for correct completion of critical steps and (4) remains functional under a variety of perturbations [16]. Dysregulation of the cell cycle control system is associated with multiple diseases, such as cancer and neurodegenerative disorders. [17]. The complex nature of cell cycle regulation with multiple interconnected and nested feedback loops combined with the known challenges of fitting the parameters of oscillatory systems to experimental data [18] exacerbate the development of mechanistically detailed, yet comprehensive cell cycle models. Nevertheless, we here sought to leverage recent technological advancements to improve on existing mammalian cell cycle models, especially with regard to four dimensions: model structure, validation data, validation methodology and model reusability. We approached the challenges of this endeavour by breaking down the problem in achievable subproblems. First, we developed models of individual cell cycle transitions. Second, we fused the cell cycle transition submodels to a model of the full cell cycle. Third, we reconstructed time courses of cell cycle regulators from highly multiplexed single cell measurement. Finally we cast this data and the cell cycle model into an optimisation problem and used a powerful metaheuristic to fit the model to the data. In this manner we aggregated existing knowledge about mammalian cell cycle dynamics to an executable model that explains the dynamics of 16 observables in human retinal pigment epithelial-1 (RPE-1) cells. By combining abstract model definition in the BioNetGen (BNG) language [14, 19] with an intuitive naming convention using Human Genome Organization Gene Nomenclature Committee (HGNC) short names, and by concretely describing the model, data and optimisation problem in the PEtab format [15], we released the model in an easily contributable form to GitHub (https://github.com/paulflang/cellcycle_petab/).

## Results

### The restriction point submodel

We start with a description of our cell cycle model, which consists of submodels for the restriction point (RP), the G1/S transition, the G2/M transition and the metaphase/anaphase (M/A) transition. For the precise meaning of abbreviations for modelled molecular species, please refer to Supplementary Table 1. The RP is a checkpoint in G1 that prevents cell cycle progression, until it is lifted by continuous or pulse like mitogen signalling [20], which stimulates cyclin D expression. The RP is defined as a point after which progression through the cell cycle no longer requires mitogen signalling [21]. Here, we use cyclin D:Cdk4/6 complex (CycD) as a proxy for mitogen signalling. The molecular basis of the restriction point is described by the Rb/E2f pathway shown in Fig. 1a [22, 23]. E2f is a transcription factor that drives the transcription of cyclin E. Before the restriction point, E2f is kept inactive by binding to Rb [24]. However, this binding is inhibited by phosphorylation of Rb, which is mediated, for example, by CycD and CycE. The ordinary differential equations (ODEs) describing the Rb/E2F pathway are listed in Supplementary Equations 1. Even without E2f autoactivation (i.e. replacing the ODE for tE2f with *tE*2*f* = 0.5; Fig. S1a, b) the mutual inhibition of CycE and Rb, combined with the inhibitor ultrasensitivity conferred by Rb allows for bistability in the Rb/E2f pathway. The bistable range and thus the robustness of this network with respect to variations in CycD concentration is further increased by transcriptional autoactivation of E2f (Fig. 1b). In agreement with the notion that once the restriction point is passed, cell cycle progression becomes independent of mitogen signalling [21], flipping the toggle switch to the high CycE state is a truly irreversible process in our submodel. Reverting the system would require negative CycD concentrations. The time course simulated with CycD just above the upper bifurcation point shows how Rb, CycE and E2f approach the high CycE steady state (Fig. S1c). The increasing concentrations in CycE and E2f will trigger CycA accumulation by switching the G1/S toggle. Their values in the high CycE steady state will therefore set upper boundaries for the upper bifurcation point in the G1/S submodel.

**Figure 1.**
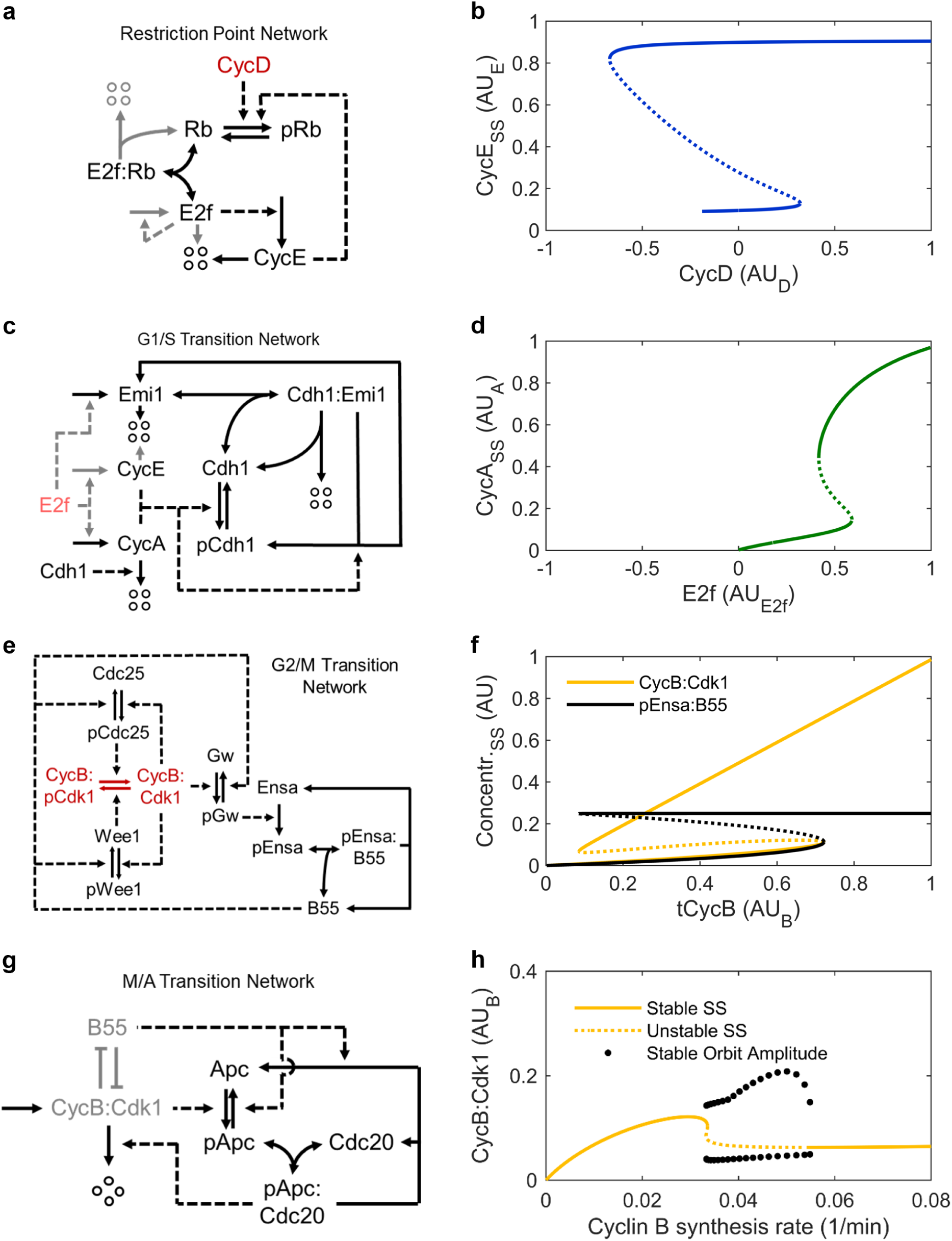
Models of individual cell cycle transitions. **a**, **b** Reaction network and bifurcation diagram of the restriction point. Lighter colour indicate reactions omitted in Fig. S1a, b. **c**, **d** Reaction network and bifurcation diagram of the G1/S transition. Lighter colour indicate reactions omitted in Fig. S2b. e, f Reaction network and bifurcation diagram of the G2/M transition as developed by Vinod and Novak [25]. **g**, **h** Reaction network and bifurcation diagram of the combined G2/M and M/A transition. Lighter colour represents a simplified schematic of the G2/M transition network shown in (e). In the reaction network diagrams conversions are represented by full arrows and catalytic interactions by dashed arrows. Four circles in5dicate degraded proteins. Red letters represent the species used as bifurcation parameters. In the bifurcation diagrams solid lines show stable and dotted lines unstable steady states. Line endings within the axes limits indicate disappearance of a steady state. For variable abbreviations please refer to Supplementary Table 1. Cdh1 and Cdh1:Emi in the G1/S transition network correspond to (p)Apc:Cdh1 and (p)Apc:Cdh1:Emi1, respectively in the full cell cycle model. Models are available in the [/versions/v0.0.1/ directory](https://github.com/paulflang/cell_cycle_model/tree/b0a820288bc98d62333acf8434174b6446df0574/versions/v0.0.1) of the cell cycle model GitHub repository.

### G1/S transition submodel

Like cyclin E, transcription of cyclin A is also activated by E2f. However, there is a delay between the accumulation of CycE and CycA proteins that shall be captured by the G1/S transition submodel. In contrast to cyclin E, cyclin A is susceptible to polyubiquitination by the Apc:Cdh1 complex. The high Apc:Cdh1 levels observed in G1 keep the CycA concentration relatively low, while the CycE concentration already rises. However, CycE and CycA can phosphorylate Cdh1, thereby preventing its association with Apc [26, 27]. Additionally, E2f also activates the transcription of Emi1 [28], which is a very slowly polyubiquitinated pseudosubstrate of Apc:Cdh1. Tight binding of Emi1 to Apc:Cdh1 further inhibits Apc:Cdh1 [29] (Fig. 1c). This ODEs for this G1/S transition network are described in Supplementary Equations 2. As total Apc appears to be in excess of total Cdh1 and tCdc20 [30], we assume in this submodel that all Cdh1 is bound to Apc. Similar G1/S transition submodels have been proposed by Barr et al. [31] and Novak and Tyson [32]. The mutual inhibition between CycA and Apc:Cdh1, combined with the inhibitor ultrasensitivity conferred by Emi1 allows for bistability in this G1/S transition network. This is illustrated in Fig. 1c, using E2f as bifurcation parameter. Setting E2f to 0.7 AU_E2F_ (i.e. just below its upper steady state in the RP submodel) and using CycE as bifurcation parameter instead, Fig. S2b shows that CycA can robustly maintain its high steady state independently of CycE. This is a critical feature of the G1/S toggle switch, since the rising concentration of CycA after flipping the switch will lead to CycE phosphorylation, marking it for SCF mediated polyubiquitination. Rising CycA concentrations will facilitate cyclin B synthesis when merging the G1/S toggle with the G2/M toggle.

### G2/M transition submodel

Cyclin B associates with Cdk1, which is kept inactive via Wee1 kinase mediated phosphorylation. Wee1 activity is counteracted by the phosphatase Cdc25. As Cdc25 is activated and Wee1 inactivated by CycB:Cdk1, this kinase phosphatase system contains two positive feedback loops. Modelling the phosphorylation and/or dephosphorylation of Cyclin B:Cdk1 with Michaelis-Menten functions as ultrasensitive Goldbeter-Koshland switch, the system can become bistable with the steady state depending on tCycB. However, recent studies suggest that the bistability of the G2/M transition network (Fig. 1 and Supplementary Equations 3) depends on the PP2A:B55/Ensa/Greatwall pathway [33, 34]. This evidence supports an earlier model by Vinod and Novak [25], which we will use to describe the G2/M transition. According to this model, the inhibitor ultrasensitivity conferred by phosphorylated Ensa does not allow for bistability if only combined with the mutual inhibition between B55 and phosphorylated Ensa. Yet, robust bistable behaviour can be observed if this inhibitor ultrasensitivity is combined with the mutual inhibition between B55 and pGw, and/or B55 and CycB:Cdk1 (Fig. 1f and Fig. S3). One critical property of the network is that the bifurcation parameter tCycB is an upper boundary of the system variable CycB:Cdk1 but not of B55. When reducing tCycB below the upper bifurcation point B55 remains bound to phosphorylated Ensa, while CycB:Cdk1 almost linearly decreases with tCycB. As B55 inactivity can stabilise its inactive state almost independently of CycB:Cdk1, it is possible to reduce tCycB to very low levels without activating B55. This feature will be exploited to reset the high tCycB at the G2/M transition to basal levels via the negative feedback loop represented by the M/A transition submodel.

### M/A transition submodel

The major hallmarks of the M/A transition are chromosome segregation and cyclin B degradation. Both processes are mediated by pApc:Cdc20, which is kept inactive via association to mitotic checkpoint proteins until all chromosomes are correctly attached to the microtubule spindle apparatus. The phosphorylation of Apc requires a shift in the ratio between Apc kinase and phosphatase activity. While it is well established that CycB:Cdk1 serves as Apc kinase [35], less is known about phosphatases acting on Apc. It has been put forth that B55 is one of the phosphatases for Cdk1 substrates [36]. In contrast to CycB:Cdk1, pEnsa:B55 at the upper bifurcation point is much lower than the pEnsa:B55 at the lower bifurcation point (Fig. 1f), allowing to couple the bistable G2/M switch with B55 driven negative feedback (Fig. 1g). Combining the negative feedback from the MA transition submodel (Supplementary Equations 4) with the G2/M switch (Supplementary Equations 3) results in limit cycle oscillations (Fig. 1h, Fig. S4) in a similar way to what has already been shown by Ferrell [37]. However, these oscillations disappear when cyclin B synthesis decreases below 0.03335 min*^−^*^1^ (Fig. 1i). This feature will be exploited in the full cell cycle model, where cyclin B synthesis will be reduced at the M/A transition to keep its concentration low in G1 phase.

### The core cell cycle model

Next, we fused the four submodels to a stably oscillating cell cycle model (Fig. 2a, b) in a stepwise manner, accounting for additional species and reactions created through the fusion (Supplementary Note 1). This core model also retained a fully functional CycD controllable restriction point (Fig. 2c, d). Without changing parameters, we could further demonstrate that the model also agrees with knockout experiments of CycE [38, 39] or CycA [40] (Fig. 2e, f).

**Figure 2.**
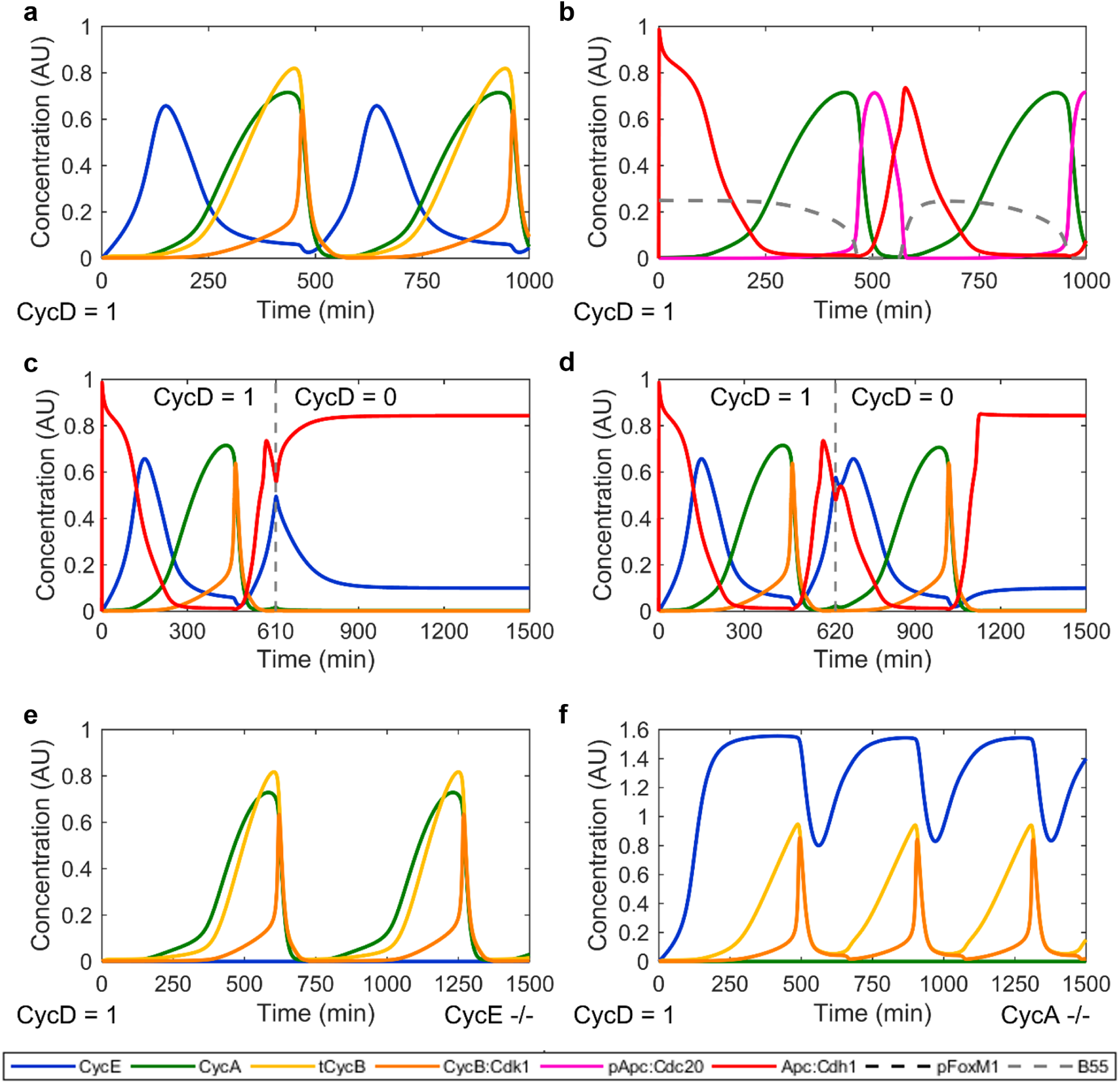
Time courses of the core cell cycle model. **a**, **b** Core cell cycle model at CycD = 1 AU_D_. **c**, **d** Simulation of mitogen deprivation. The full cell core model was simulated as in (a, b) until 610 min (c) and 620 min (d), respectively. CycD as proxy for mitogen availability was then turned to 0 AU_D_ and the simulation was continued until 1500 min. **e** Full cell cycle model with CycE knockout. **f** Full cell cycle model with CycA knockout. For variable abbreviations please refer to Supplementary Table 1. The model is available in the [/versions/v1.0.0/directory](https://github.com/paulflang/cell_cycle_model/tree/b0a820288bc98d62333acf8434174b6446df0574/versions/v1.0.0) of the cell cycle model GitHub repository.

### Implementing a DNA damage checkpoint

After minor model refinements and rescaling (Changelogs and model versions in Supplementary Table 2; simulation comparison in Fig. S5a), we aimed at extending the model by a DNA damage checkpoint. However, due to the complexity of the model, extending the model in its ODE-based form would be a painstaking and error prone task. Therefore, we decided to work on a more abstract level and converted the ODE-based model to a rule-based model defined in the BioNetGen (BNG) language [14, 19] (Fig. S5b). Using the BNG syntax, we further introduced a systematic naming convention of all molecular species, based on Human Genome Organization Gene Nomenclature Committee (HGNC) short names and known important phosphorylation sites. This naming convention is explained in Supplementary Note 2 and the mapping to the names used in the above models is shown in Supplementary Table 1. For instance, protein A with an unoccupied binding side for protein B, an occupied binding site C and a phosphorylated residue Res123 would be written as A(B, C!+, Res123∼p). The representation of the model in the BNG language facilitated introduction of a DNA damage checkpoint through inhibition of cyclins via binding of unphosphorylated cyclin kinase inhibitor CDKN1A (also known as p21). CDKN1A expression is controlled via TP53, a transcription factor that is activated by DNA damage. Polyubiquitination by the SKP2-containing SCF complex marks CDKN1A for degradation. Noteworthy, the mutual inhibition of CDKN1A and Cdks, combined with the ultrasensitivity conferring stoichiometric binding, results in bistability of the DNA damage checkpoint [41]. After adding CDKN1A, TP53, SKP2 and the rules coarsely described in words above, TP53 provides an interface for activating a DNA damage response checkpoint (version 3.1.0). Fig. 3 shows TP53 induced reversible activation of DNA damage checkpoints in G1 and G2 phase. Closer inspection of the G1 checkpoint indicates that CDKN1A is not fully reduced to baseline by the time the cell cycle continues, as indicated by rising CCNA/B levels. We also observe high levels of the proliferative effector CCNE before CCNA/B levels rise, which may titrate some CDKN1A away from CCNA/B. These simulation results are in good agreement with Stallaert et al. [42], who found that the re-entry trajectory does not follow the same path as the exit trajectory. Instead, they argue that the arrest state is not reversed, but overcome through elevated expression of proliferative effectors. However, any interpretation of our simulation results must be treated with caution, as the simulation was performed with arbitrarily chosen parameters and arbitrary units of concentration. To further increase robustness of the bistability at the G1/S transition, we added CDKN1B as another stoichiometric inhibitor of cyclins, leading to version 3.2.0 (Supplementary Note 3).

**Figure 3.**
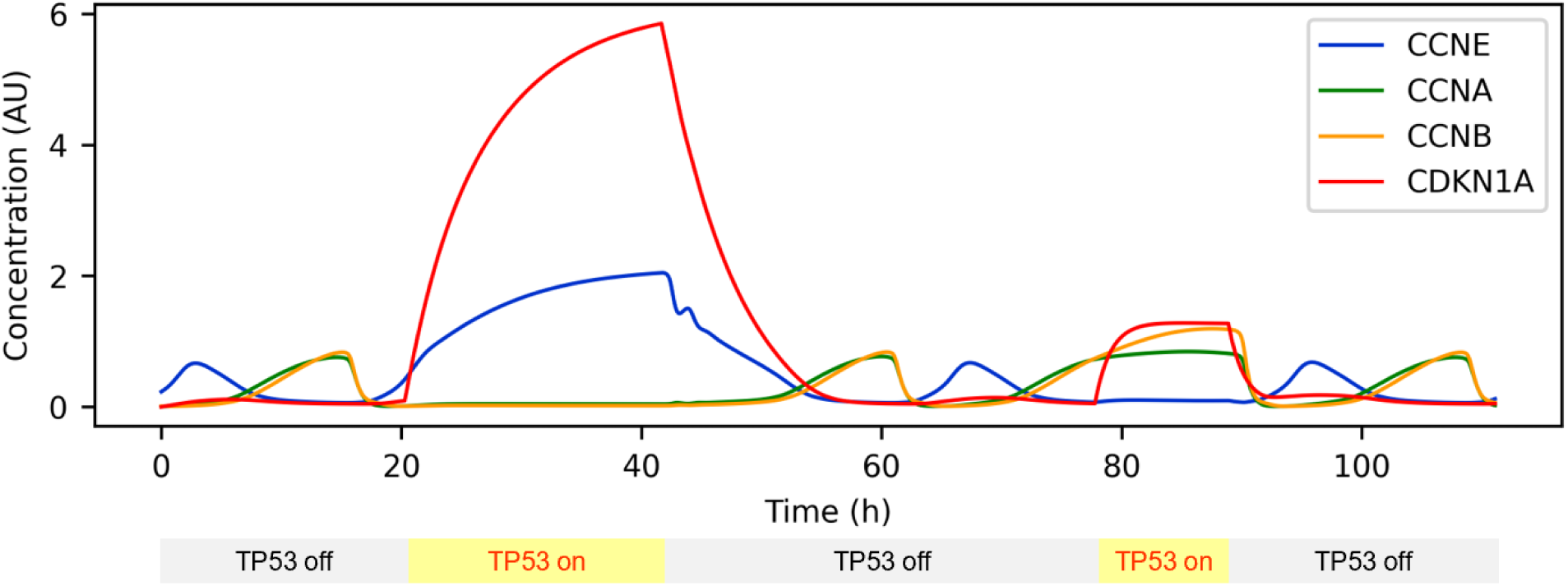
Time courses with alternating TP53 levels. DNA damage was simulated by activating TP53. The first activation corresponds to G1 phase, the second to G2 phase. Inactivation of TP53 allows continued proliferation. For better visualisation the G1 checkpoint was lifted before the steady state was reached. Model version 3.1.0.

### Introducing the notion of compartmentalisation

The model presented so far assumes uniform spatial distribution of the cell cycle regulators. However, there is substantial evidence of nucleocytoplasmic shuttling of several cell cycle regulators [43–46], including evidence that such translocation plays important roles in cell cycle regulation, for instance as a source of bistability [47]. Such nucleocytoplasmic shuttling through nuclear pores is carried out by the Ran-GTP cycle, which depends on nuclear export and import signals on the transported proteins, which can be (de)activated by posttranslational modifications [48]. To enable more accurate representations of cell cycle control, we implemented the capacity for nucleocytoplasmic shuttling of cell cycle regulators into the model (Supplementary Note 4). This capability will later enable parameter optimisation algorithms to tune import/export rate constants. If identifiable, these rate constants may point towards translocation mechanisms of hard-to-observe species, such as complexes and certain phosphoproteins. We confirmed sustained cell cycle oscillations by simulating for more than 100 doubling times (Fig. S6). In addition, we implemented nuclear envelope breakdown (Fig. S7). However, this led to significantly reduced simulation speed. Therefore, nuclear envelope breakdown was switched off for parameter estimation.

### Obtaining multiplexed and spatially resolved time course measurements of cell cycle regulators

To estimate some of the 325 unknown parameters in the compartmental model of the cell cycle, we were searching for highly multiplexed and spatially resolved time course measurements of cell cycle regulators. We found that Stallaert and colleagues [42] published single-cell snapshot measurements of cell cycle regulators in asynchronously dividing human retinal pigment epithelial-1 (RPE-1) cells. These measurements were obtained from iterative indirect immunofluorescence imaging (4i) experiments [49]. To reconstruct pseudotime courses from such single-cell measurements, several cell trajectory reconstruction algorithms have been developed over the last decade [6–11]. For the particular case of reconstructing cell cycle trajectories, we can (a) exploit that the proliferative cell cycle trajectory forms a closed circle and (b) cell division creates two cells from one. More precisely, from Kafri *et al.* [6] we know that the cell density *p* at cell age *t* of a perfectly asynchronous cell population moving across a single circular trajectory can be described as

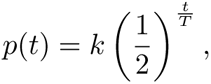

where *k* is a scaling constant and *T* the doubling time of the cells. We also know that the whole population size *N* is the area under this curve

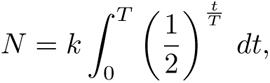

and therefore

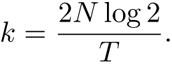

The number *r* of cells younger than *t* is therefore

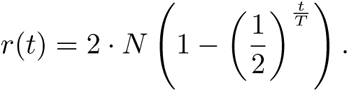

Rearranging we obtain the cell cycle time from *r* as

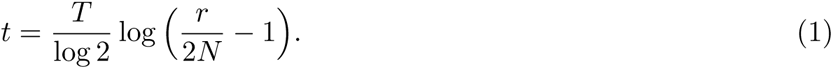

### Testing the reCAT trajectory reconstruction algorithm on simulated data

To test cell cycle trajectory reconstruction with reCAT, 300 cells were sampled from model version 2.1.4, such that cell density decreases exponentially over the cell cycle with one cell cycle length half-life. First, all 49 model variables were provided for trajectory reconstruction without adding noise methods on data transformation. reCAT was able to perfectly reconstruct the cell cycle trajectory from this dataset (Person correlation coefficient: *R* = 1.0). As a next step, the input data was corrupted with *lognormal*(1, 0.8^2^) noise. Despite the substantial noise level, reCAT recovered the original cell cycle trajectory with near perfect correlation (*R* = 0.997). This accuracy remained practically unchanged (*R* = 0.987) even after reducing the information content of the data by eliminating 40 variables from the reconstruction procedure (Fig. S8).

### Reconstructing time courses from imaging data

Stallaert *et al.* [42] obtained spatially resolved measurements of 48 cell cycle regulators in thousands of RPE-1 cells. Using the dimensionality reduction method PHATE (Potential of Heat-diffusion for Affinity-based Trajectory Embedding) [50], they found that a significant proportion of RPE-1 cells exit the proliferative cell cycle trajectory into a non-proliferative G0 arm. We discarded such G0 cells prior to trajectory reconstruction (Fig. S9). For reasons outlined in Supplementary Note 5, we believe that discarding G0 cells is sufficient to safely apply Equation (1) to calculate cell cycle pseudo-times from ranks. After removal of such G0 cells, we performed trajectory reconstruction on 300 randomly selected cells (Methods). reCAT was only provided with the 36 of the 40 most predictive cell cycle features identified by Stallaert *et al.* that did not contain missing values. The other features were ignored during reconstruction, as they may contribute more noise than cell cycle information. Ranking the cells resulted in time courses of all 292 features, 12 of which are shown in Fig. 4 (the full dataset is available in table 4i_stallaert/2021_Stallaert_cycling_reCAT/smoothened_data.xlsx of the cell cycle time course GitHub repository, and Kalman-smoothened data can be inspected with [this interactive figure](https://paulflang.github.io/cell_cycle_time_course/4i_stallaert/2021_Stallaert_cycling_reCAT/kalman.html)). Overall, the reconstructed cell cycle trajectory corresponded well with known behaviour of cell cycle regulators. For instance, CCNA accumulated in later cell cycle stages and was abruptly degraded in mitosis. CCNB followed the same pattern with a slight delay and DNA content doubled throughout the cell cycle. Interestingly however, CCNE concentration appeared more stable across the cell cycle compared to life cell imaging experiments in HeLa cells [31] and immunofluorescence measurements in U2OS cells [51].

**Figure 4.**
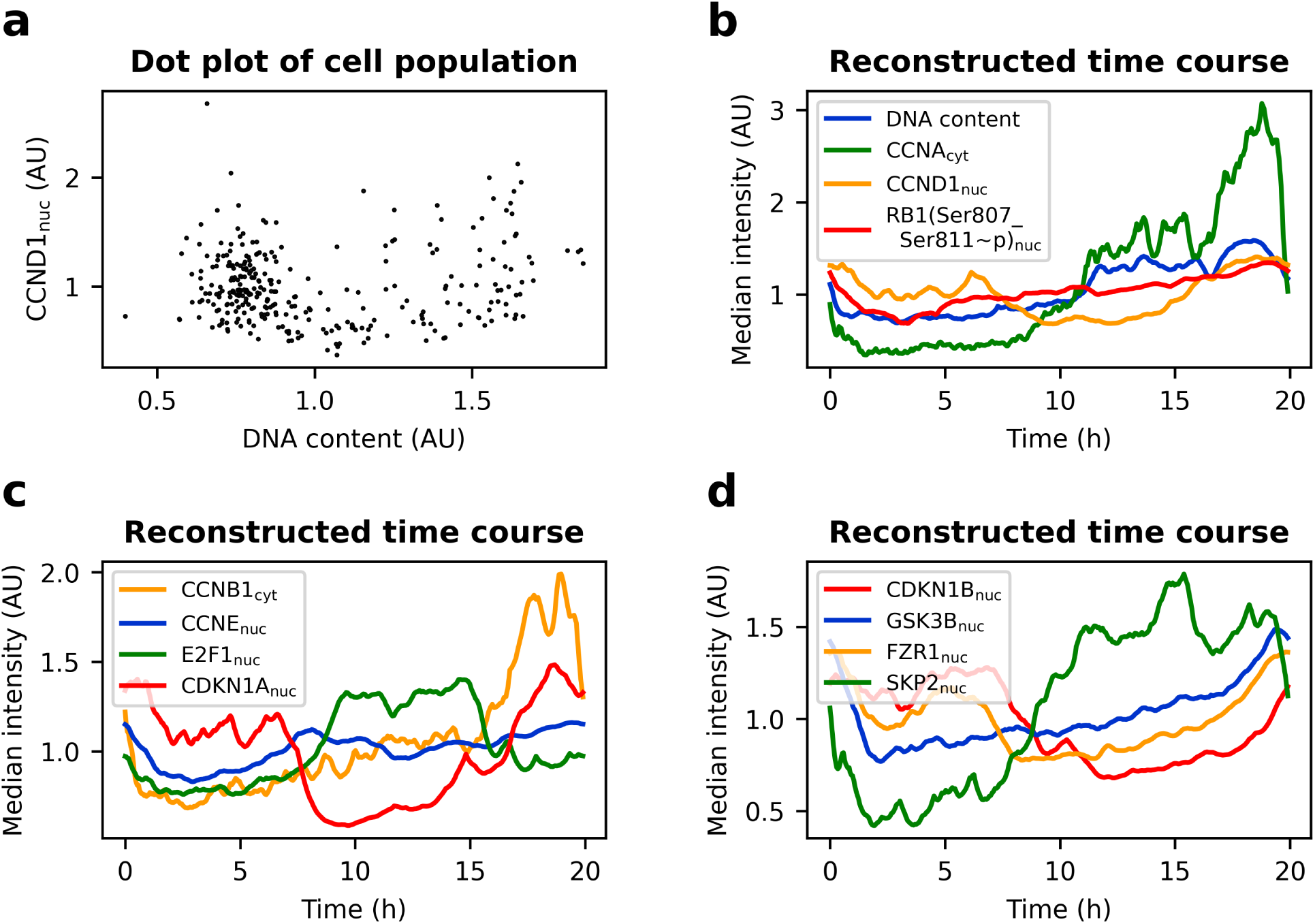
Time course of cell cycle regulators in RPE-1 cells. Proliferating cells of publicly available 4i measurements of an asynchronously dividing RPE-1 population [42] were gated in PHATE space (Fig. S9). For better visibility the variables are normalised to a mean of one, and axes are clipped. **a** Dot plot of 300 untreated RPE-1 cells. **b-d** Reconstructed time course of the cell population from (a). reCAT was provided with 36 variables. Time was calculated using Equation (1) and data was smoothened using a Kalman filter.

### Parameter estimation with perfect simulated data

Having developed a compartmental model of cell cycle control and reconstructed spatially resolved time courses of cell cycle regulators, we aimed to estimate the parameters of our model. However, the optimization problem at hand involved several challenges, including: unknown parameter bounds, high dimensional parameter space, unknown parameters in the observation function that maps model states to fluorescent readout, unknown contributions to and magnitude of measurement noise, presence of multiple local optima, and structural and practical parameter unidentifiabilities. Managing these challenges is still a matter of ongoing research [18, 52–55]. To nevertheless identify an algorithm that is capable of finding the globally optimal parameters of the cell cycle model, we first performed the optimisation on idealized simulated data. These data were generated by simulating one cell cycle and sampling 100 time points for all states without introducing noise. Using multistart optimization provided with gradients obtained from adjoint-sensitivity analysis, we found 100 different and poor solutions from 100 starting points, indicating a highly rugged objective function. To avoid such premature attraction to local optima, we therefore switched to Cooperative enhanced Scatter Search (CeSS), a hybrid global-local optimisation algorithm [56] (Methods) that performed well in benchmark tests [57]. Optimisation took 10.7 hours using four cores of a personal computer (Intel® CoreTM i7-8550U CPU @ 1.80GHz). This time, the ground truth parameters used for generating the data were recovered with only minor deviations, except for the initial conditions of Apc:Cdh and pCdh (Fig. 5a). In the cell cycle model total Cdh is constant. CeSS has recovered the correct amount for *Apc* : *Cdh*(*t* = 0) + *pCdh*(*t* = 0) = 0.87 up to the second decimal digit, but estimated the initial condition of Apc:Cdh higher and pCdh lower than their respective ground truth values. As the observation function was simply the pure species abundance, this confusion of initial values increases the objective function value. Therefore, the otherwise near perfect recovery of ground truth parameters indicates structural identifiability of the posed problem. Had there been structural unidentifiabilities, that is perfectly flat valleys in the objective function, it would be a highly unlikely coincidence that CeSS returns the ground truth parameters instead of any other point in the valley. Fig. 5b shows the practically perfect overlap between the time course of the noise-free simulated datapoints and the simulation with estimated parameters.

**Figure 5.**
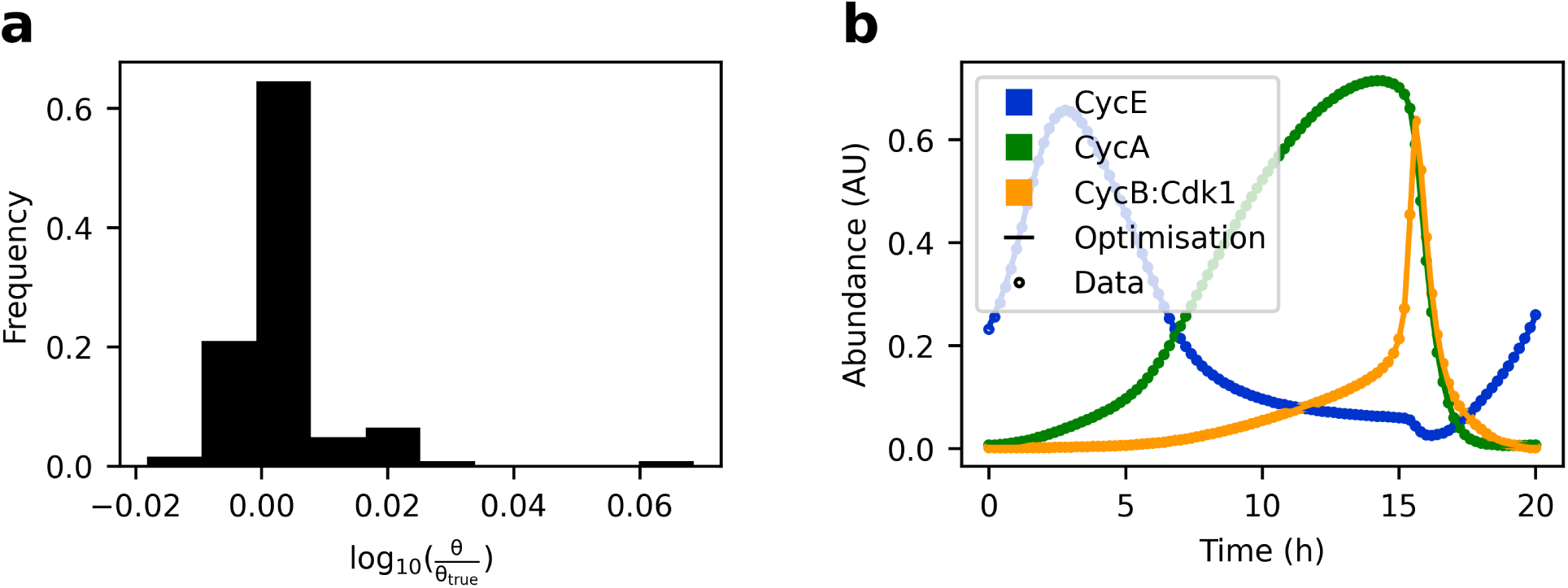
Testing Cooperative enhanced Scatter Search (CeSS). 101 evenly spaced, simulated, noise-free datapoints of 41 variables were generated with model version 2.0.0 (circles). Dynamic parameters were estimated within a [0.1 *·* ***θ****_true_,* 10 *·* ***θ****_true_*] search window. Initial conditions were estimated within a [*eps,* 1.5 *·* ***θ****_true_*] search window (except for Apc, where the upper boundary was 5). **a** Histogram showing distribution of all parameters *θ* versus ground truth except for *Apc* : *Cdh* and *pCdh* (see text), and *kDpE*2*f* 1 and *kPhC*25*A* (*θ_true_* = 0 and *θ ≤ eps*). **b** Simulated time courses for three of the 41 variables, using the parameter set *θ* found by CeSS. *eps*: machine epsilon.

### Imitating real imaging data and speeding up optimisation

Having found CeSS as an optimizer that is capable of solving the cell cycle optimisation problem on idealized data using a personal computer, the next step was to speed up optimisation to meet the demands of realistic data. To this end we used self-adaptive cooperative enhanced scatter search (saCeSS). saCeSS improves on CeSS by successive adaptation of hyperparameters, improving the cooperation strategy and timing, and combining fine- and coarse-grained parallelisation on high performance computing hardware [58]. The version used for this work is available in the sacess_cell_cycle_petab Bitbucket repository. To test saCeSS on model version 3.0.0, the optimisation problem was updated to better mimic our experimental data. In particular, the Stallaert *et al.* dataset includes five observables that are informative for this version of the cell cycle problem: CCNE, CCNA, CCNB1, E2F1 and RB1(Ser807 Ser811∼p). The expected density of the 300 datapoints is exponentially decreasing over the cell cycle, with a half-life of one doubling time. Importantly, the observables represent antibody fluorescent intensities and were rescaled to a mean of one. This means that observables *y* map to model species via

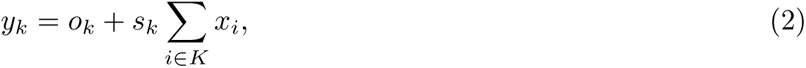

where the index *k* denotes different antibodies, *o_k_* represents an offset constant, *s_k_* a scaling constant, *K* the set of all species to which the k^th^ antibody binds, and *x* their concentration. The resulting optimisation problem including the species-to-observable mapping from Equation (2), observable noise, simulated data, and estimated parameters with bounds was specified in the PEtab format and can be found in the [/versions/v3.0.0/PEtab_PL_v3_0_0sim/](https://github.com/paulflang/cell_cycle_petab/tree/d562d3d4d19921ff851c06d0854b8b3f05c63d85/versions/v3.0.0/PEtab_PL_v3_0_0sim) directory of the cell cycle petab GitHub repository. The simulated observables were corrupted with *N* (1, 0.1^2^) multiplicative noise and *o_k_* was set to zero for all *k*. Since several species are no longer directly observed and scaling constants are introduced as new parameters, the problem is no longer expected to be structurally identifiable. To nevertheless find good fits to the data, two different local solvers were tested for the optimisation: parPE (ipopt solver using adjoint sensitivity gradients) [55], and gradient-free dynamic hill climbing (DHC). PEtab problems specify the objective as posterior probability. Here, uniform priors and a standard deviation for the measurement noise of 0.1 were used, rendering the problem equivalent to least-squares optimisation constrained to parameter bounds. Both solvers went under the objective function value of the ground truth parameters, indicating slight overfitting. As expected, neither solver recovered the ground truth parameters, which can be attributed to overfitting and parameter unidentifiabilities. The lowest objective function value was found with the parPE local solver option in saCeSS. A simulation with the corresponding parameter values and the convergence curve is shown in Fig. S10.

### Parameter estimation with imaging data

Next, we exchanged the artificial data with real experimental data from Stallaert *et al*. This required us to make five modifications to the optimisation strategy (Supplementary Note 6). In particular, we had to fit over two cycles to enforce oscillatory behaviour. All these strategies were combined into a new problem specification, using the parameters from the oscillatory but non-fitted model as initial guess. The best performance was achieved with the DHC local solver option in saCeSS. The found solution led to simulations that are in good agreement with experimentally observed time courses. Fig. S11 shows the convergence curve and directly compares nuclear measurements and the simulation.

The optimisation procedure was repeated for model version 3.2.0, which contains relevant variables to use three additional nuclear observables in the Stallaert [59] dataset: SKP2, CDKN1A and CDKN1B. The corresponding PEtab problem can be found in the /versions/v3.2.0/ directory of the cell cycle petab GitHub repository. Fig. S12 compares the simulation results using the estimated parameters to experimental measurements.

Finally, we estimated the parameters of model version 4.0.0 using nuclear and cytoplasmic observables. Since the measurement noise in the cytoplasm was much lower than in the nucleus, we allowed observable-specific noise by adding the following noise formula to the optimisation problem:

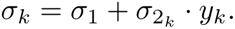

*σ*_1_ and *σ*_2_*k* were estimated as part of the optimisation problem and represent additive and observable-specific multiplicative noise, respectively. *y_k_* denotes the k^th^ observable and *σ_k_* the standard deviation of normally-distributed measurement noise. The corresponding PEtab problem can be found in the [/versions/v4.0.0/](https://github.com/paulflang/cell_cycle_petab/tree/d562d3d4d19921ff851c06d0854b8b3f05c63d85/versions/v4.0.0) directory of the cell cycle petab GitHub repository. Even for this compartmental model of the cell cycle, saCeSS found a solution that leads to simulations that are in excellent agreement with experimentally observed location and time courses of the measured cell cycle regulators (Fig. 6). While this solution was initially located at an edge of the search window, shifting the search window and optimising for another 40 h did not further improve the solution.

**Figure 6.**
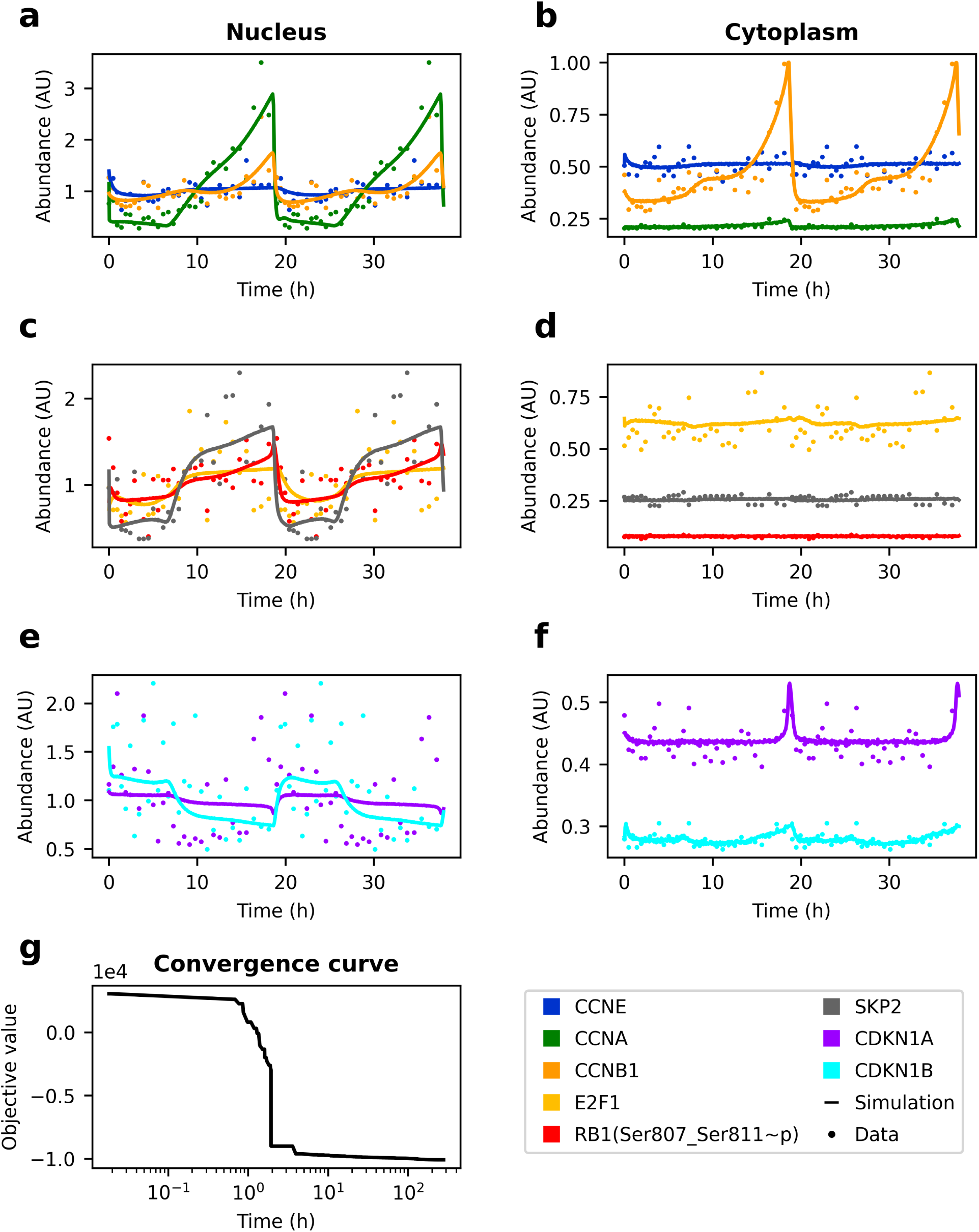
Model version 4.0.0 fitted to pseudo-time courses of RPE-1 cells. Experimental measurements and simulation results using estimated parameters. For better visibility, only every 10^th^ measurement is shown. **a, c, e** Nuclear compartment. **b, d, f** Cytoplasmic compartment. **g** Convergence curve.

## Discussion

Cell cycle control is one of the most central mechanisms of life and dysregulation in mammals results in various diseases such as cancer and neurodegenerative diseases [17]. Generating and improving computational models of the cell cycle has therefore been a long-standing goal of systems biology [60–67]. With the present work, we contribute the most complete mammalian cell cycle model known to the authors that accurately explains time courses of experimentally determined observables. In particular, we improved on comparable existing models (Table 1) along the four dimensions of model structure (e.g. using the notion of compartmentalisation and DNA checkpoints in the same model), validation data (i.e. using the largest number of densely sampled observables), validation methodology (i.e. using a probabilistically motivated objective function and a more reproducible optimisation procedure) and reusability (i.e. providing the model and optimisation problem in multiple community standards, including an easily extensible rule-based description, tracking significant parts of model development with git version control, exploiting BNGL syntax and HGNC short names for concise semantics and releasing the model to the interactive GitHub platform). Yet, the parameters of the model were structurally unidentifiable and predictions for small molecule or genetic cell cycle perturbations could not be tested in lack of appropriate experimental time course data. Generating such data will thus be the next significant step towards improving and evaluating the model. In addition, we invite the community to contribute suggestions and improvements of any form via GitHub issues and pull requests.

**Table 1.**
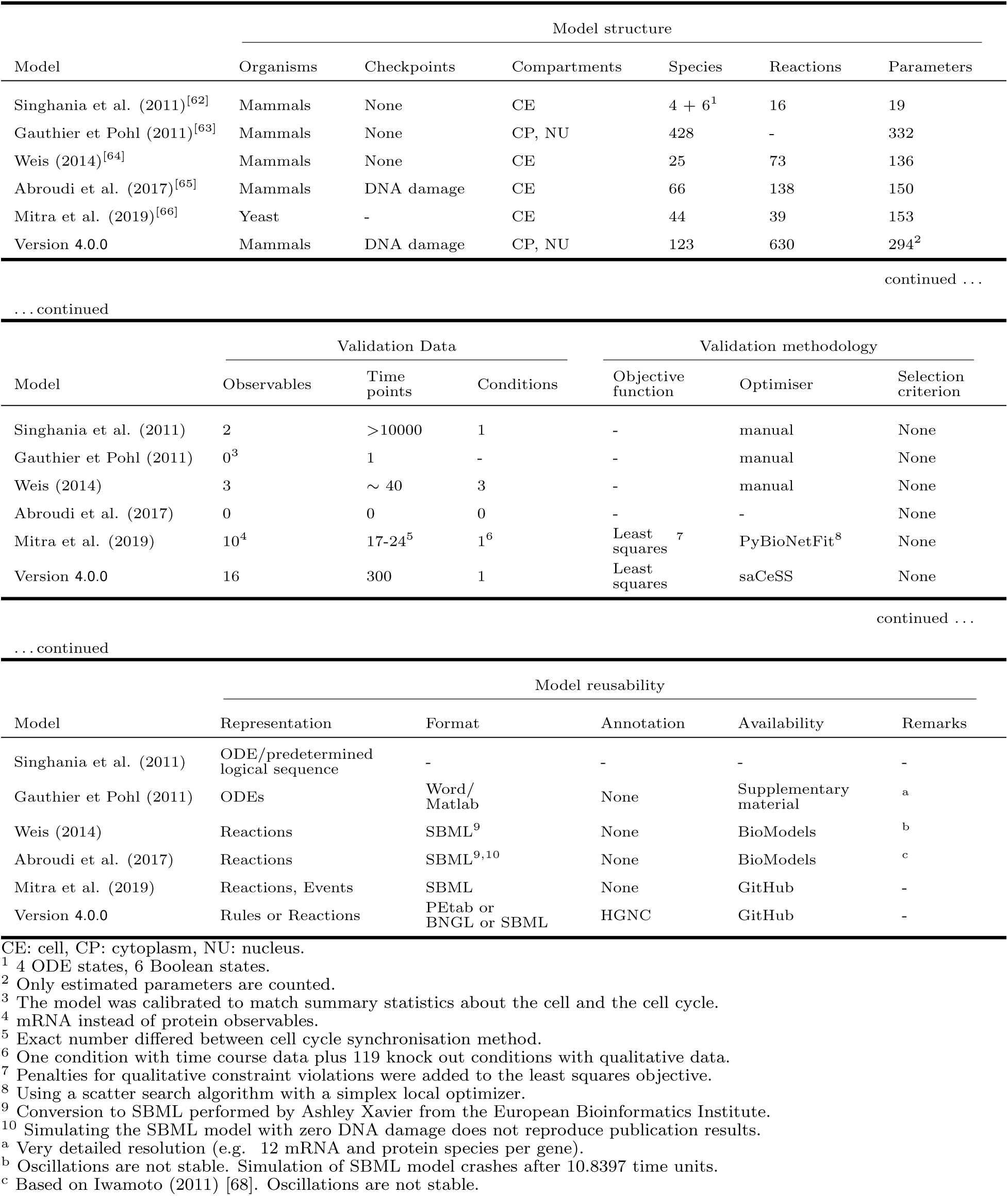
Comparison of this work with existing cell cycle models.

| Model | Model structure |  |  |  |  |  |
| --- | --- | --- | --- | --- | --- | --- |
|  | Organisms | Checkpoints | Compartments | Species | Reactions | Parameters |
| Singhanian et al. (2011)<sup>[62]</sup> | Mammals | None | CE | 4 + 6<sup>1</sup> | 16 | 19 |
| Gauthier et Pohl (2011)<sup>[63]</sup> | Mammals | None | CP, NU | 428 | - | 332 |
| Weis (2014)<sup>[64]</sup> | Mammals | None | CE | 25 | 73 | 136 |
| Abroudi et al. (2017)<sup>[65]</sup> | Mammals | DNA damage | CE | 66 | 138 | 150 |
| Mitra et al. (2019)<sup>[66]</sup> | Yeast | - | CE | 44 | 39 | 153 |
| Version 4.0.0 | Mammals | DNA damage | CP, NU | 123 | 630 | 294<sup>2</sup> |

| Model | Validation Data |  |  | Validation methodology |  |  |
| --- | --- | --- | --- | --- | --- | --- |
|  | Observables | Time points | Conditions | Objective function | Optimiser | Selection criterion |
| Singhanian et al. (2011) | 2 | >10000 | 1 | - | manual | None |
| Gauthier et Pohl (2011) | 0<sup>3</sup> | 1 | - | - | manual | None |
| Weis (2014) | 3 | ~ 40 | 3 | - | manual | None |
| Abroudi et al. (2017) | 0 | 0 | 0 | - | - | None |
| Mitra et al. (2019) | 10<sup>4</sup> | 17-24<sup>5</sup> | 1<sup>6</sup> | Least squares<sup>7</sup> | PyBioNetFit<sup>8</sup> | None |
| Version 4.0.0 | 16 | 300 | 1 | Least squares | saCeSS | None |

| Model | Model reusability |  |  |  |  |
| --- | --- | --- | --- | --- | --- |
|  | Representation | Format | Annotation | Availability | Remarks |
| Singhanian et al. (2011) | ODE/predetermined logical sequence | - | - | - | - |
| Gauthier et Pohl (2011) | ODEs | Word/ Matlab | None | Supplementary material | <sup>a</sup> |
| Weis (2014) | Reactions | SBML<sup>9</sup> | None | BioModels | <sup>b</sup> |
| Abroudi et al. (2017) | Reactions | SBML<sup>9,10</sup> | None | BioModels | <sup>c</sup> |
| Mitra et al. (2019) | Reactions, Events | SBML | None | GitHub | - |
| Version 4.0.0 | Rules or Reactions | PETab or BNGL or SBML | HGNC | GitHub | - |

CE: cell, CP: cytoplasm, NU: nucleus.
<sup>1</sup> 4 ODE states, 6 Boolean states.<sup>2</sup> Only estimated parameters are counted.<sup>3</sup> The model was calibrated to match summary statistics about the cell and the cell cycle.<sup>4</sup> mRNA instead of protein observables.<sup>5</sup> Exact number differed between cell cycle synchronisation method.<sup>6</sup> One condition with time course data plus 119 knock out conditions with qualitative data.<sup>7</sup> Penalties for qualitative constraint violations were added to the least squares objective.<sup>8</sup> Using a scatter search algorithm with a simplex local optimizer.<sup>9</sup> Conversion to SBML performed by Ashley Xavier from the European Bioinformatics Institute.<sup>10</sup> Simulating the SBML model with zero DNA damage does not reproduce publication results.<sup>a</sup> Very detailed resolution (e.g. 12 mRNA and protein species per gene).<sup>b</sup> Oscillations are not stable. Simulation of SBML model crashes after 10.8397 time units.<sup>c</sup> Based on Iwamoto (2011) [68]. Oscillations are not stable.

## Methods

### Developing the Core model

The core model and the transition models are formulated as systems of ODEs. Nullcline plots, bifurcation diagrams and time courses where calculated with the freely available software XPP/XPPAUT [69]. For steady state bifurcation analysis the bifurcation parameter was set to zero and XPP/XPPAUT was started from the corresponding steady state. This steady state bifurcation analysis also detects Hopf bifurcations. Periodic bifurcation analysis (Fig. 1h) was started from the Hopf bifurcation at *k_SyCb_ ≈* 0.055 min^-1^. The biochemical reactions making up the RP, G1/S and G2M submodels were chosen to comply with experimental observations of bistability [22, 31, 70–72] in these networks. These submodels where merged in a stepwise manner to yield an RP-G1S, RP-G1S-G2M, G2M-MA and finally a full cell cycle model (RP-G1S-G2M-MA). Additional reactions and species where added whenever necessary and to make the model more comprehensive. Reaction parameters were chosen based on Williams *et al.* [73], where applicable. For the remaining parameters, an attempt was made to fulfil the following criteria: (a) Parameters shall be biologically reasonable compared to the parameters obtained by Williams *et al.*. (b) Parameters in the RP, G1/S and G2M submodels shall allow for bistability. (c) After merging the submodels to the full model, parameters were adjusted to obtain sustained oscillations as observed in unperturbed cells, CycE [38] and CycA [40] knockouts to obtain model version 1.0.0.

### Including DNA damage response and compartmentalisation

Prior to interfacing with DNA damage response regulator and compartmentalisation, the model underwent rescaling, minor changes and conversion to BioNetGen Language version 2.5.2 [19] using RuleBender version 2.3.1 [74] (Supplementary Table 2, versions 1.0.0–3.0.0). The precise changes after model version 3.0.0, including the implementation of DNA damage response and compartmentalisation were documented and tracked with Git version control and made available on the cell_cycle_model GitHub repository.

### Model verification

All relevant changes to the reaction network and all conversions between model description formats were subject to model verification. Version 1.0.0 was checked by rewriting the ODEs in a chemical reaction-based format using the freely available software COPASI [75]. The chemical reactions were automatically converted to a system of ODEs that were used for manual proofreading of the XPP/XPPAUT code. Time course simulations of XPP/XPPAUT and COPASI lead to identical results within a small margin of tolerance for numerical inaccuracy. Similarly, conversion from chemical reactions (version 2.1.4) to reaction rules (version 3.0.0) lead to identical simulation results within a small margin of tolerance. Starting with version 3.0.0, for all modifications to the reaction rules and molecule types, the expected new number of reactions generated by each rule and the expected number of total species were documented in files /versions/v*/sim_log_expected.log)^1^ of the cell_cycle_model GitHub repository. Equivalency with the auto-generated number of reactions per rule and total species (files /cell_cycle_model/versions/v*/sim_log.log) was confirmed by manually comparing relevant differences with the Visual Studio Code (Microsoft) file comparison tool.

### Major model assumptions

Major assumptions include: (1) All reactions (even non-elementary reactions) are best modelled with mass action kinetics (exception: Hill function as phenomenological description of nuclear pore phosphorylation by CCNB(CDK1_Thr14_Tyr15∼u); see Supplementary Table 1 and Supplementary Note 2 for the naming convention used). (2) Cyclin:Cdk complex concentration is proportional to cyclin concentration. (3) The time for protein expression is negligibly small. (4) The model environment provides all protein subunits, metabolites, components of the transcription-translation and protein degradation machinery that are required by the reactions described in the model. It also provides constant total levels of CCND(), RB1(), FZR1(), ENSA_ARPP19(), PPP2R2B(), MASTL(), WEE1(), CDC25(), APC(), TP53() and DNA (see Supplementary Table 1 for alternative names of these proteins). (5) (De)phosphorylation of a protein is not changed by binding to another protein (exceptions: PPP2R2B(ENSA_ARPP19) actively dephosphorylates ENSA_ARPP19(PPP2R2B!?, Ser62_Ser67∼p).FZR1(nTerm∼u) phosphorylation is in the full model assumed to be reduced due to putative steric hindrances within the APC(FZR!1,FBXO5!2).FZR1(APC!1,FBXO5!3,nTerm∼u).FBXO5(APC!2,FZR1!3) complex).

(6) PPP2R2B (ENSA_ARPP19) is a putative APC phosphatase. (7) Phosphorylated cyclin E and A are immediately degraded. (8) Effects of local enrichment of the modelled components can be ignored.

### Cleaning of experimental data

Outlier cells with respect to the following metrics were excluded from further analyses of the publicly available data from Stallaert *et al.* [59]: nuclear area, DNA content, CDKN1A(Thr145∼p), cellular area, cytoplasmic DAPI stain in all rounds, and ratio between cellular and nuclear area (indicating segmentation errors). Exact thresholds are available in the curation directories of the respective datasets on the cell_cycle_time_course GitHub repository. Values are displayed by hovering the mouse over the dashed red line indicating the threshold. Figures can be opened as described in the repository README.

### Trajectory reconstruction with reCAT

reCAT was provided with 300 cells sampled across one cell cycle model simulation (version 2.1.4). Variables were transformed according to

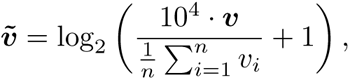

where ***v*** denotes a variable and *n* is the number of cells. reCAT was run with default settings resulting in an ordered list of cells in a cycle. The direction and start of the cycle were chosen automatically from patterns of cell cycle variables and corrected manually if necessary. For the Stallaert dataset manual correction was informed by manual annotation of cell age and phase. Raw (non-transformed) variables were divided by their mean. A Kalman filter was applied to the time series, using the R programming language with functions StructTS and tsSmooth. The time series was padded with the last 10% of the cell cycle at the beginning and the first 10% of the cell cycle in the end. StructTS optimised the maximum likelihood of the ”level” model, where the evolution of variable *x*(*t*) is described by

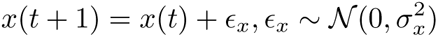

and the measurement *y*(*t*) by

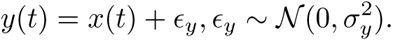

Paddings were removed and the order of the cells was converted to pseudo-time via Equation (1).

### Multistart optimisation and CeSS

Optimisation problems were specified and optimised as described by the [software supplement](https://zenodo.org/record/1304034#.YmBXvdqZOHs) of Villaverde *et al* [57]. For CeSS, the following hyperparameters were chosen: opts.ndiverse=’auto’; opts.maxeval=5e6; opts.log var=[]; opts.local.solver=’fmincon’; opts.local.finish=’fmincon’; opts.local.bestx=0; opts.local.tol=1; opts.local.n1=100; opts.dim refset=’auto’; opts.prob bound=0.5;

opts.n threads=8; opts.n iter=10; opts.maxtime cess=1*3600. The hyperparameters for threads 1–8 were set to dim1=3; bal1=0; n2 1=4; dim2=5; bal2=0.25; n2 2=7; dim3=5; bal3=0.25; n2 3=10; dim4=7.5; bal4=0.25; n2 4=15; dim5=10; bal5=0.5; n2 5=20; dim6=12; bal6=0.25; n2 6=50; dim7=15; bal7=0.25; n2 7=100; dim8=15; bal8=0.5; n2 8=10000000000. AMICI was used for model simulations and gradient calculations [76].

### Optimisation with saCeSS

All problems were specified in the PEtab format and can be found in the cell_cycle_petab GitHub repository. saCeSS was interfaced with parPE and used AMICI for model simulations and gradient calculations (code available in the sacess_cell_cycle_petab Bitbucket repository). Default settings of saCeSS and parPE were used (saCeSS hyperparameters are automatically adapted based on the number of islands). The number of runs (coarse-grained parallelisation) and processors (fine-grained parallelisation) is documented in /versions/PEtab_PLang_problem_v43.xlsx of the cell cycle petab GitHub repository.

### Data visualisation and illustrations

Data were visualised using MATLAB (R2018a) and Python (v3.9.5 and v3.10.3) with the matplotlib (v3.4.3 and v3.5.1) package. For plotting Fig. S6 the first cell cycle was eliminated from the simulation results as its initial conditions do not match the conditions at the start of subsequent cycles. Interactive online figures were were created with the plotly library (v4.9.1) in R (v3.6.1). Reaction Networks were drawn with Microsoft PowerPoint.

## Acknowledgements

The authors would like to thank Frank Stefan Heldt, John Sekar and Jonathan Karr for inspiring the conceptualisation of this work.

## Footnotes

1 The asterisk is a placeholder for the version name. For v3.0.1, /versions/v3.0.1/n species reactions expected.txt was used instead of a .log file.

## Notes

https://github.com/paulflang/cell_cycle_petab

### Competing Interest Statement

The authors declare that the model is protected by an Academic Use License (https://github.com/paulflang/cell cycle petab/blob/main/LICENCE) and not for use by commercial businesses, unless licensed by Oxford University Innovation Ltd.

### Summary of Updates

Added supplementary information.

## References

[1] Fuchou Tang et al. “mRNA-Seq whole-transcriptome analysis of a single cell”. Nature Methods 6.5 (2009), pp. 377–382.

[2] Dmitry R. Bandura et al. “Mass Cytometry: Technique for Real Time Single Cell Multitarget Immunoassay Based on Inductively Coupled Plasma Time-of-Flight Mass Spectrometry”. Analytical Chemistry 81.16 (2009), pp. 6813–6822.

[3] Bogdan Budnik et al. “SCoPE-MS: mass spectrometry of single mammalian cells quantifies proteome heterogeneity during cell differentiation”. Genome Biology 19.1 (2018), p. 161.

[4] Anne E. Carpenter et al. “CellProfiler: image analysis software for identifying and quantifying cell phenotypes”. Genome Biology 7.10 (2006), R100.

[5] Laurens van der Maaten and Geoffrey Hinton. “Visualizing Data using t-SNE”. Journal of Machine Learning Research 9.86 (2008), pp. 2579–2605.

[6] Ran Kafri et al. “Dynamics extracted from fixed cells reveal feedback linking cell growth to cell cycle”. Nature 494.7438 (2013), pp. 480–483.

[7] Gabriele Gut et al. “Trajectories of cell-cycle progression from fixed cell populations”. Nature Methods 12.10 (2015), pp. 951–954.

[8] Laleh Haghverdi et al. “Diffusion pseudotime robustly reconstructs lineage branching”. Nature Methods 13.10 (2016), pp. 845–848.

[9] Zehua Liu et al. “Reconstructing cell cycle pseudo time-series via single-cell transcriptome data”. Nature Communications 8.1 (2017), p. 22.

[10] Maria Anna Rapsomaniki et al. “CellCycleTRACER accounts for cell cycle and volume in mass cytometry data”. Nature Communications 9.1 (2018), p. 632.

[11] Luca Rappez et al. “DeepCycle reconstructs a cyclic cell cycle trajectory from unsegmented cell images using convolutional neural networks”. Molecular Systems Biology 16.10 (2020), e9474.

[12] M. Hucka et al. “The systems biology markup language (SBML): a medium for representation and exchange of biochemical network models”. *Bioinformatics (Oxford*, England) 19.4 (2003), pp. 524–531.

[13] Autumn A. Cuellar et al. “An Overview of CellML 1.1, a Biological Model Description Language”. SIMULATION 79.12 (2003), pp. 740–747.

[14] Michael L. Blinov et al. “BioNetGen: software for rule-based modeling of signal transduction based on the interactions of molecular domains”. Bioinformatics 20.17 (2004), pp. 3289–3291.

[15] Leonard Schmiester et al. “PEtab—Interoperable specification of parameter estimation problems in systems biology”. PLOS Computational Biology 17.1 (2021), e1008646.

[16] John J. Tyson and Bela Novak. “Temporal Organization of the Cell Cycle”. Current Biology 18.17 (2008), R759–R768.

[17] B. Zhivotovsky and S. Orrenius. “Cell cycle and cell death in disease: past, present and future”. Journal of Internal Medicine 268.5 (2010), pp. 395–409.

[18] Jake Alan Pitt and Julio R. Banga. “Parameter estimation in models of biological oscillators: an automated regularised estimation approach”. BMC Bioinformatics 20.1 (2019), p. 82.

[19] Leonard A. Harris et al. “BioNetGen 2.2: advances in rule-based modeling”. Bioinformatics 32.21 (2016), pp. 3366–3368.

[20] Y. Zwang et al. “Two phases of mitogenic signaling unveil roles for p53 and EGR1 in elimination of inconsistent growth signals”. Mol Cell 42.4 (2011), pp. 524–35.

[21] A. B. Pardee. “A restriction point for control of normal animal cell proliferation”. Proc Natl Acad Sci USA 71.4 (1974), pp. 1286–90.

[22] G. Yao et al. “A bistable Rb-E2F switch underlies the restriction point”. Nat Cell Biol 10.4 (2008), pp. 476–82.

[23] Frank S. Heldt et al. “A comprehensive model for the proliferation-quiescence decision in response to endogenous DNA damage in human cells”. Proceedings of the National Academy of Sciences of the United States of America 115.10 (2018), pp. 2532–2537.

[24] J. William Harbour and Douglas C. Dean. “The Rb/E2F pathway: expanding roles and emerging paradigms”. Genes & Development 14.19 (2000), pp. 2393–2409.

[25] P. K. Vinod and Bela Novak. “Model scenarios for switch-like mitotic transitions”. FEBS Letters 589.6 (2015), pp. 667–671.

[26] Wolfgang Zachariae et al. “Control of Cyclin Ubiquitination by CDK-Regulated Binding of Hct1 to the Anaphase Promoting Complex”. Science 282.5394 (1998), pp. 1721–1724.

[27] Alan W. Lau et al. “Regulation of APCCdh1 E3 ligase activity by the Fbw7/cyclin E signaling axis contributes to the tumor suppressor function of Fbw7”. Cell Research 23.7 (2013), pp. 947–961.

[28] J. Y. Hsu et al. “E2F-dependent accumulation of hEmi1 regulates S phase entry by inhibiting APC(Cdh1)”. Nat Cell Biol 4.5 (2002), pp. 358–66.

[29] J. J. Miller et al. “Emi1 stably binds and inhibits the anaphase-promoting complex/cyclosome as a pseudosubstrate inhibitor”. Genes Dev 20.17 (2006), pp. 2410–20.

[30] J. Y. Huang and J. W. Raff. “The dynamic localisation of the Drosophila APC/C: evidence for the existence of multiple complexes that perform distinct functions and are differentially localised”. J Cell Sci 115.Pt 14 (2002), pp. 2847–56.

[31] Alexis R. Barr et al. “A Dynamical Framework for the All-or-None G1/S Transition”. Cell Systems 2.1 (2016), pp. 27–37.

[32] Béla Novák and John J. Tyson. “Mechanisms of signalling-memory governing progression through the eukaryotic cell cycle”. Current Opinion in Cell Biology. Cell Signalling 69 (2021), pp. 7–16.

[33] Scott Rata et al. “Two Interlinked Bistable Switches Govern Mitotic Control in Mammalian Cells”. Current Biology 28.23 (2018), 3824–3832.e6.

[34] Nadia Hégarat, et al. “Cyclin A triggers Mitosis either via the Greatwall kinase pathway or Cyclin B”. The EMBO Journal 39.11 (2020), e104419.

[35] Bruce Alberts et al. “Chapter 17: The Cell Cycle”. Molecular Biology of the Cell. Taylor & Francis Group, 2014.

[36] S. Mochida and T. Hunt. “Protein phosphatases and their regulation in the control of mitosis”. EMBO Rep 13.3 (2012), pp. 197–203.

[37] J. E. Ferrell. “Feedback loops and reciprocal regulation: recurring motifs in the systems biology of the cell cycle”. Curr Opin Cell Biol 25.6 (2013).

[38] S. Ortega et al. “Cyclin-dependent kinase 2 is essential for meiosis but not for mitotic cell division in mice”. Nat Genet 35.1 (2003), pp. 25–31.

[39] Y. Geng et al. “Cyclin E ablation in the mouse”. Cell 114.4 (2003), pp. 431–43.

[40] I. Kalaszczynska et al. “Cyclin A - Redundant in Fibroblasts, Essential in Hematopoietic and Embryonal Stem Cells”. Cell 138.2 (2009), pp. 352–65.

[41] K. Wesley Overton et al. “Basal p21 controls population heterogeneity in cycling and quiescent cell cycle states”. Proceedings of the National Academy of Sciences 111.41 (2014), E4386–E4393.

[42] Wayne Stallaert et al. “The structure of the human cell cycle”. Cell Systems 0.0 (2021).

[43] Anja Hagting et al. “Translocation of cyclin B1 to the nucleus at prophase requires a phosphorylation-dependent nuclear import signal”. Current Biology 9.13 (1999), pp. 680–689.

[44] Andrew B. Gladden and J. Alan Diehl. “Location, location, location: The role of cyclin D1 nuclear localization in cancer”. Journal of Cellular Biochemistry 96.5 (2005), pp. 906–913.

[45] Iordanka A. Ivanova, Alisa Vespa, and Lina Dagnino. “A Novel Mechanism of E2F1 Regulation Via Nucleocytoplasmic Shuttling: Determinants of Nuclear Import and Export”. Cell Cycle 6.17 (2007), pp. 2186–2195.

[46] Helena Silva Cascales et al. “Cyclin A2 localises in the cytoplasm at the S/G2 transition to activate PLK1”. Life Science Alliance 4.3 (2021).

[47] Silvia D. M. Santos et al. “Spatial Positive Feedback at the Onset of Mitosis”. Cell 149.7 (2012), pp. 1500–1513.

[48] Tommaso Cavazza and Isabelle Vernos. “The RanGTP Pathway: From Nucleo-Cytoplasmic Transport to Spindle Assembly and Beyond”. Frontiers in Cell and Developmental Biology 3 (2016).

[49] Gabriele Gut, Markus D. Herrmann, and Lucas Pelkmans. “Multiplexed protein maps link subcellular organization to cellular states”. Science 361.6401 (2018), eaar7042.

[50] Kevin R. Moon et al. “Visualizing structure and transitions in high-dimensional biological data”. Nature Biotechnology 37.12 (2019), pp. 1482–1492.

[51] Diana Mahdessian et al. “Spatiotemporal dissection of the cell cycle with single-cell proteogenomics”. Nature 590.7847 (2021), pp. 649–654.

[52] A. Raue et al. “Structural and practical identifiability analysis of partially observed dynamical models by exploiting the profile likelihood”. Bioinformatics 25.15 (2009), pp. 1923–1929.

[53] Alejandro F. Villaverde, Nikolaos Tsiantis, and Julio R. Banga. “Full observability and estimation of unknown inputs, states and parameters of nonlinear biological models”. Journal of The Royal Society Interface 16.156 (2019), p. 20190043.

[54] Fabian Frõhlich, Carolin Loos, and Jan Hasenauer. “Scalable Inference of Ordinary Differential Equation Models of Biochemical Processes”. Gene Regulatory Networks: Methods and Protocols. Methods in Molecular Biology. Springer, 2019, pp. 385–422.

[55] Leonard Schmiester et al. “Efficient parameterization of large-scale dynamic models based on relative measurements”. Bioinformatics 36.2 (2020), pp. 594–602.

[56] Alejandro F. Villaverde, Jose A. Egea, and Julio R. Banga. “A cooperative strategy for parameter estimation in large scale systems biology models”. BMC Systems Biology 6.1 (2012), p. 75.

[57] Alejandro F. Villaverde et al. “Benchmarking optimization methods for parameter estimation in large kinetic models”. Bioinformatics 35.5 (2019), pp. 830–838.

[58] David R. Penas et al. “Parameter estimation in large-scale systems biology models: a parallel and self-adaptive cooperative strategy”. BMC Bioinformatics 18.1 (2017), p. 52.

[59] Wayne Stallaert, et al. “The structure of the human cell cycle”. bioRxiv (2021), p. 2021.02.11.430845.

[60] B. Stern and P. Nurse. “A quantitative model for the cdc2 control of S phase and mitosis in fission yeast”. Trends Genet 12.9 (1996), pp. 345–50.

[61] Katherine C. Chen et al. “Kinetic Analysis of a Molecular Model of the Budding Yeast Cell Cycle”. Molecular Biology of the Cell 11.1 (2000), pp. 369–391.

[62] R. Singhania et al. “A hybrid model of mammalian cell cycle regulation”. PLoS Comput Biol 7.2 (2011), e1001077.

[63] John H Gauthier and Phillip I Pohl. “A general framework for modeling growth and division of mammalian cells”. BMC Systems Biology 5.1 (2011), p. 3.

[64] Michael C. Weis et al. “A Data-Driven, Mathematical Model of Mammalian Cell Cycle Regulation”. PLOS ONE 9.5 (2014), e97130.

[65] Ali Abroudi, Sandhya Samarasinghe, and Don Kulasiri. “A comprehensive complex systems approach to the study and analysis of mammalian cell cycle control system in the presence of DNA damage stress”. Journal of Theoretical Biology 429 (2017), pp. 204–228.

[66] Eshan D. Mitra et al. “PyBioNetFit and the Biological Property Specification Language”. iScience 19 (2019), pp. 1012–1036.

[67] Ulrike Münzner, Edda Klipp, and Marcus Krantz. “A comprehensive, mechanistically detailed, and executable model of the cell division cycle in Saccharomyces cerevisiae”. Nature Communications 10.1 (2019), p. 1308.

[68] Kazunari Iwamoto et al. “Mathematical modeling of cell cycle regulation in response to DNA damage: Exploring mechanisms of cell-fate determination”. Biosystems 103.3 (2011), pp. 384–391.

[69] Bard Ermentrout. *Simulating, Analyzing, and Animating Dynamical Systems*. Software, Environments and Tools. Society for Industrial and Applied Mathematics, 2002.

[70] W. Sha et al. “Hysteresis drives cell-cycle transitions in Xenopus laevis egg extracts”. Proc Natl Acad Sci USA 100.3 (2003), pp. 975–80.

[71] Joseph R. Pomerening, Eduardo D. Sontag, and James E. Ferrell Jr. “Building a cell cycle oscillator: hysteresis and bistability in the activation of Cdc2”. Nature Cell Biology 5.4 (2003), pp. 346–351.

[72] Alexis R. Barr et al. “DNA damage during S-phase mediates the proliferation-quiescence decision in the subsequent G1 via p21 expression”. Nature Communications 8 (2017), p. 14728.

[73] Byron C Williams et al. “Greatwall-phosphorylated Endosulfine is both an inhibitor and a substrate of PP2A-B55 heterotrimers”. eLife 3 (2014), e01695.

[74] Adam M. Smith et al. “RuleBender: integrated modeling, simulation and visualization for rule-based intracellular biochemistry”. BMC Bioinformatics 13.8 (2012), S3.

[75] S. Hoops et al. “COPASI–a COmplex PAthway SImulator”. Bioinformatics 22.24 (2006), pp. 3067–74.

[76] Fabian Frõhlich, et al. “AMICI: high-performance sensitivity analysis for large ordinary differential equation models”. Bioinformatics 37.20 (2021), pp. 3676–3677.

