## Supplementary Information for "Reusable rule-based cell cycle model explains compartment-resolved dynamics of 16 observables in RPE-1 cells"

**Table 1** — Variables of the core model.

| ODE variable (Unit[1]) | Description | BNGL equivalent |
|---|---|---|
| t (min) | time | time |
| CycE ($\mathrm{AU_{E}}$) | cyclin E:Cdk2 complex | `CCNE()` |
| tE2f ($\mathrm{AU_{E2f}}$) | total transcription factor E2f | `E2F(DBD!?,RB1!?,Ser332)` |
| E2f ($\mathrm{AU_{E2f}}$) | free, unphosphorylated E2f | `E2F(DBD,RB1,Ser332~u)` |
| pE2f ($\mathrm{AU_{E2f}}$) | free, phosphorylated E2f | `E2F(DBD,RB1,Ser332~p)` |
| Rb ($\mathrm{AU_{E2f}}$) | unphosphorylated retinoblastoma protein (incl. (p)E2f:Rb) | `RB1(E2F!?,Ser807_Ser811~u)` |
| pRb ($\mathrm{AU_{E2f}}$) | phosphorylated retinoblastoma protein (always free) | `RB1(E2F,Ser807_Ser811~p)` |
| fRb ($\mathrm{AU_{E2f}}$) | free, unphosphorylated Rb | `RB1(E2F,Ser807_Ser811~u)` |
| pE2f:Rb ($\mathrm{AU_{E2f}}$) | phosphorylated E2f bound to retinoblastoma protein | `E2F(DBD,RB1!1,Ser332~p).`<br>`RB1(E2F!1,Ser807_Ser811~u)` |
| E2f:PX ($\mathrm{AU_{Px}}$) | E2f bound to the promoter of gene X (CycA, CycE, E2f, FoxM1) | `E2F(DBD!1,RB1,Ser332~u).`<br>`*_promoter(E2F!1)` |
| E2f:Rb ($\mathrm{AU_{E2F}}$) | unphosphorylated E2f bound to retinoblastoma protein | `E2F(DBD,RB1!1,Ser332~u).`<br>`RB1(E2F!1,Ser807_Ser811~u)` |
| CycA ($\mathrm{AU_{A}}$) | cyclin A:Cdk1/2 complex | `CCNA()` |
| tEmi1 ($\mathrm{AU_{Cdh1}}$) | total early mitotic inhibitor 1 (Emi1) | `FBXO5(APC!?,FZR1,Ser182)` |
| Emi1 ($\mathrm{AU_{Cdh1}}$) | free, unphosphorylated Emi1 | `FBXO5(APC,FZR1,Ser182~u)` |
| pEmi1 ($\mathrm{AU_{Cdh1}}$) | phosphorylated Emi1 (always free) | `FBXO5(APC,FZR1,Ser182~p)` |
| Apc ($\mathrm{AU_{Cdh1}}$) | unphosphorylated anaphase promoting complex (APC/C) without substrate adaptor subunit | `APC(FZR1_CDC20,FBXO5,Ser355~u)` |
| pApc ($\mathrm{AU_{Cdh1}}$) | free, phosphorylated APC/C | `APC(FZR1_CDC20,FBXO5,Ser355~p)` |
| Cdh1 ($\mathrm{AU_{Cdh1}}$) | free, unphosphorylated APC/C substrate adaptor subunit Cdh1 | `FZR1(APC,FBXO5,nTerm~u)` |
| pCdh1 ($\mathrm{AU_{Cdh1}}$) | phosphorylated Cdh1 (always fee) | `FZR1(APC,FBXO5,nTerm~p)` |
| Apc:Cdh1 ($\mathrm{AU_{Cdh1}}$) | APC/C bound to Cdh1 | `APC(FZR1_CDC20!1,FBXO5,Ser355~u).`<br>`FZR1(APC!1,FBXO5,nTerm~u)` |
| pApc:Cdh1 ($\mathrm{AU_{Cdh1}}$) | phosphorylated APC/C bound to Cdh1 | `APC(FZR1_CDC20!1,FBXO5,Ser355~p).`<br>`FZR1(APC!1,FBXO5,nTerm~u)` |
| Apc:Cdh1:Emi1 ($\mathrm{AU_{Cdh1}}$) | APC/C bound to Cdh1 and Emi1 | `APC(FZR1_CDC20!1,FBXO5!2,Ser355~u).`<br>`FZR1(APC!1,FBXO5!3,nTerm~u).`<br>`FBXO5(APC!2,FZR1!3,Ser182~u)` |
| pApc:Cdh1:Emi1 ($\mathrm{AU_{Cdh1}}$) | phosphorylated APC/C bound to Cdh1 and Emi1 | `APC(FZR1_CDC20!1,FBXO5!2,Ser355~p).`<br>`FZR1(APC!1,FBXO5!3,nTerm~u).`<br>`FBXO5(APC!2,FZR1!3,Ser182~u)` |
| tFoxM1 ($\mathrm{AU_{Fox}}$) | total transcription factor FoxM1 | `FOXM1(DBD!?,Thr600)` |
| pFoxM1 ($\mathrm{AU_{Fox}}$) | phosphorylated (i.e. active) transcription factor FoxM1 | `FOXM1(DBD,Thr600~p)` |
| tCycB ($\mathrm{AU_{B}}$) | cyclin B:(p)Cdk1 complex | `CCNB(CDK1_Thr14_Tyr15)` |
| pFoxM1:PCycB ($\mathrm{AU_{Pb}}$) | phosphorylated FoxM1 bound to the promoter of cyclin B | `FOXM1(DBD!1,Thr600~p).`<br>`CCNB_promoter(FOXM1!1)` |
| CycB:Cdk1 ($\mathrm{AU_{B}}$) | cyclin B in complex with unphosphorylated (i.e. active) Cdk1 | `CCNB(CDK1_Thr14_Tyr15~u)` |
| Wee1 ($\mathrm{AU_{Wee}}$) | unphosphorylated (i.e. active) Wee1 kinase | `WEE1(Ser123~u)` |
| pCdc25 ($\mathrm{AU_{C25}}$) | phosphorylated (i.e. active) Cdc25 phosphatase | `CDC25(pSites~p)` |
| pGw ($\mathrm{AU_{Gw}}$) | phosphorylated Greatwall kinase | `MASTL(Thr198~p)` |
| pEnsa ($\mathrm{AU_{Ensa}}$) | total (incl. pEnsa:B55) phosphorylated forms of Endosulfine alpha and Arpp19 | `ENSA_ARPP19(PPP2R2B!?,`<br>`Ser62_Ser67~p)` |
| B55 ($\mathrm{AU_{Ensa}}$) | protein phosphatase 2A core dimer in complex with the regulatory subunit B55 | `PPP2R2B(ENSA_ARPP19)` |
| pEnsa:B55 ($\mathrm{AU_{Ensa}}$) | phosphorylated Ensa bound to B55 complex | `ENSA_ARPP19(PPP2R2B!1,`<br>`Ser62_Ser67~p).`<br>`PPP2R2B(ENSA_ARPP19!1)` |
| tCdc20 ($\mathrm{AU_{Cdh1}}$) | total APC/C substrate adaptor subunit Cdc20 | `CDC20(APC!?)` |
| pFoxM1:PCdc20 ($\mathrm{AU_{Pc}}$) | phosphorylated FoxM1 bound to the promoter of Cdc20 | `FOXM1(DBD!1,Thr600~p).`<br>`CDC20_promoter(FOXM1!1)` |
| Cdc20 ($\mathrm{AU_{Cdh1}}$) | free Cdc20 | `CDC20(APC)` |
| pApc:Cdc20 ($\mathrm{AU_{Cdh1}}$) | phosphorylated APC/C bound to Cdc20 | `APC(FZR1_CDC20!1,`<br>`FBXO5,Ser355~p).CDC20(APC!1)` |

[1] As arbitrary units (AU) of dimension "concentration" with characteristic scale indicated in subscript.
* one of CCNE, CCNA, E2F, FBXO5, FOXM1.

**Table 2** — Changelog of model versions.

| Version | Changes |
|---|---|
| 1.0.0 | - |
| 2.0.0 | - Rescaled cell cycle length to 20 h (H1 human embryonic stem cells).<br>- Hard coded/eliminated parameters f1-f4 that were once introduced to describe loss of enzymatic access if target is in complex.<br>- Renamed species to avoid ambiguity in case-insensitive software.<br>- Ran version 1.0.0 for 9950 time units and used this state as initial conditions in version 2.0.0.<br>- Deleted observables. |
| 2.1.0 | - Removed the simplifying assumption from the ODE model that (un)binding of transcription factor to promoter does not affect the concentration of free transcription factor. |
| 2.1.1 | - Created separate promoters for Ce, Ca, E2f, Emi and Fox from one E2f activated promoter (formerly called Px). |
| 2.1.2 | - Introduced observables for parameter optimisation purposes (were never used in the end and specified via PEtab instead). |
| 2.1.3 | - Introduced interfaces for small molecule mediated inhibition of Ce and transcriptional/translational inhibition for Cb (were never used in the end). |
| 2.1.4 | - Introduced interface for small molecule mediated inhibition of Wee1 and removed it for Ce. |
| 3.0.0 | - Converted model into BioNetGen language.<br>- Introduced a systematic naming convention. |
| 3.0.1 | - Make E2F binding to DNA and its transcriptional activity independent of E2F phosphorylation state.<br>- Introduce baseline APC dephosphorylation.<br>- Allow dephosphorylation of DNA-bound FOXM1. |
| 3.1.0 | - Added DNA damage checkpoint (SKP2 and TP53 and CDKN1A). |
| 3.2.0 | - Added CDKN1B. |
| 4.0.0 | - Added compartmentalisation. |

### Supplementary Equations 1: The restriction point submodel

Please refer to the /versions/v0.0.1/ directory of the cell_cycle_model GitHub repository for parameter values and initial conditions. Note that for Supplementary Equations (1)–(26) it is assumed that transcription factors are present in much higher concentration than promoters, i.e. promoter binding has negligible effect on free transcription factor concentration. This assumption will be removed for full cell cycle model versions ≥ v2.1.0 (see also Table 2)

$$\frac{dRb}{dt} = k_{DpRb} \cdot pRb - k_{PhRb} \cdot (CycD + CycE) \cdot Rb + k_{DeE2f} \cdot E2f{:}Rb \tag{1}$$

$$\frac{dCycE}{dt} = k_{SyCe1} + k_{SyCe2} \cdot E2f{:}Px - k_{DeCe} \cdot CycE \tag{2}$$

$$\begin{aligned}\frac{dE2f}{dt} &= k_{SyE2f1} + k_{SyE2f2} \cdot E2f{:}Px - k_{DeE2f} \cdot E2f \\ &\quad + k_{DiE2fRb} \cdot E2f{:}Rb - k_{AsE2fRb} \cdot E2f \cdot fRb\end{aligned} \tag{3}$$

$$\frac{dtE2f}{dt} = k_{SyE2f1} + k_{SyE2f2} \cdot E2f{:}Px - k_{DeE2f} \cdot tE2f \tag{4}$$

$$\frac{dE2f{:}Px}{dt} = k_{AsEPx} \cdot E2f \cdot (t_{Dna} - E2f{:}Px) - k_{DiEPx} \cdot E2f{:}Px \tag{5}$$

$$pRb = t_{Rb} - Rb \tag{6}$$

$$fRb = Rb - (tE2f - E2f) \tag{7}$$

$$E2f{:}Rb = tE2f - E2f \tag{8}$$

### Supplementary Equations 2: The G1/S transition submodel

$$\begin{aligned}\frac{dtEmi}{dt} &= k_{SyEmi1} + k_{SyEmi2} \cdot E2f{:}Px - k_{DeEmi1} \cdot tEmi1 \\ &- k_{DeEmi2} \cdot Apc{:}Cdh1{:}Emi1\end{aligned} \tag{9}$$

$$\begin{aligned}\frac{dCycA}{dt} &= k_{SyCa1} + k_{SyCa2} \cdot E2f{:}Px \\ &- (k_{DeCa1} + k_{DeCa2} \cdot Apc{:}Cdh1) \cdot CycA\end{aligned} \tag{10}$$

$$\frac{dCycE}{dt} = k_{SyCe1} + k_{SyCe2} \cdot E2f{:}Px - k_{DeCe} \cdot CycE \tag{11}$$

$$\begin{aligned}\frac{dApc{:}Cdh1{:}Emi1}{dt} &= k_{AsACE} \cdot Apc{:}Cdh1 \cdot (tEmi - Apc{:}Cdh1{:}Emi1) \\ &- (k_{DiACE} + k_{DeEmi1} + k_{DeEmi2} + k_{PhCdhE} \cdot CycE \\ &+ k_{PhCdhA} \cdot CycA) \cdot Apc{:}Cdh1{:}Emi1\end{aligned} \tag{12}$$

$$\begin{aligned}\frac{dpCdh1}{dt} &= (k_{PhCdhA} \cdot CycA + k_{PhCdhE} \cdot CycE) \cdot (t_{Cdh1} - pCdh1) \\ &- k_{DpCdh} \cdot pCdh1\end{aligned} \tag{13}$$

$$\frac{dE2f{:}Px}{dt} = k_{AsEPx} \cdot E2f \cdot (t_{Dna} - E2f{:}Px) - k_{DiEPx} \cdot E2f{:}Px \tag{14}$$

$$Apc{:}Cdh1 = t_{Cdh1} - pCdh1 - Apc{:}Cdh1{:}Emi1 \tag{15}$$

### Supplementary Equations 3: The G2/M transition submodel

$$\frac{dpEnsa}{dt} = k_{PhEnsa} \cdot pGw \cdot (t_{Ensa} - pEnsa) - k_{DpEnsa} \cdot pEnsa : B55 \tag{16}$$

$$\frac{dCycB{:}Cdk1}{dt} = r_{Cdc25} \cdot (t_{CycB} - CycB{:}Cdk1) - r_{Wee} \cdot CycB{:}Cdk1 \tag{17}$$

$$\begin{aligned}\frac{dpGw}{dt} &= k_{PhGw} \cdot CycB{:}Cdk1 \cdot (t_{Gw} - pGw) \\ &- (k_{DpGw1} + k_{DpGw2} \cdot B55) \cdot pGw\end{aligned} \tag{18}$$

$$\begin{aligned}\frac{dpEnsa{:}B55}{dt} &= k_{AspEB55} \cdot B55 \cdot (pEnsa - pEnsa{:}B55) \\ &- (k_{DipEB55} + k_{DpEnsa}) \cdot pEnsa{:}B55\end{aligned} \tag{19}$$

$$B55 = t_{B55} - pEnsa{:}B55 \tag{20}$$

$$r_{Wee} = k_{Wee1} + \frac{(k_{Wee2} - k_{Wee1}) \cdot k_{DpWee} \cdot B55}{k_{DpWee} \cdot B55 + k_{PhWee} \cdot CycB{:}Cdk1} \tag{21}$$

$$r_{Cdc25} = k_{Cdc25\_1} + \frac{(k_{Cdc25\_2} - k_{Cdc25\_1}) \cdot k_{PhCdc25} \cdot CycB{:}Cdk1}{k_{PhCdc25} \cdot CycB{:}Cdk1 + k_{DpCdc25} \cdot B55} \tag{22}$$

### Supplementary Equations 4: The M/A transition submodel

$$\begin{aligned}\frac{dpApc}{dt} &= k_{PhApc} \cdot (t_{Apc} - pApc) \cdot (CycB : Cdk1) \\ &- k_{DpApc} \cdot B55 \cdot pApc\end{aligned} \tag{23}$$

$$\begin{aligned}\frac{dpApc{:}Cdc20}{dt} &= k_{AsAC20} \cdot (t_{Cdc20} - pApc{:}Cdc20) \cdot (pApc - pApc{:}Cdc20) \\ &- (k_{DiAC20} + k_{DpApc} \cdot B55) \cdot pApc{:}Cdc20\end{aligned} \tag{24}$$

$$\frac{dtCycB}{dt} = k_{SyCb} - (k_{DeCb1} + k_{DeCb2} \cdot pApc{:}Cdc20) \cdot t_{CycB} \tag{25}$$

$$\begin{aligned}\frac{dCycB{:}Cdk1}{dt} &= k_{SyCb} - (k_{DeCb1} + k_{DeCb2} \cdot pApc{:}Cdc) \cdot CycB{:}Cdk1 \\ &+ r_{Cdc25} \cdot (t_{CycB} - CycB{:}Cdk1) - r_{Wee} \cdot CycB{:}Cdk1\end{aligned} \tag{26}$$

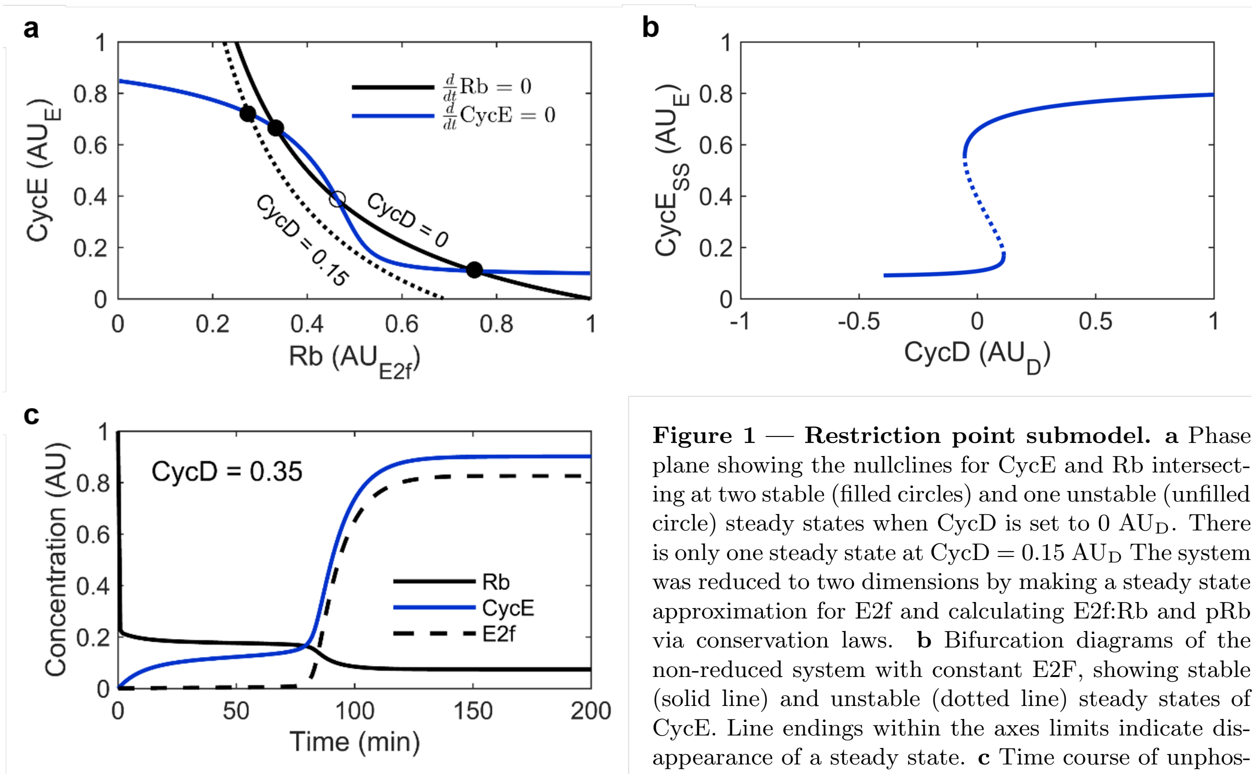


**Figure 1 — Restriction point submodel. a** Phase plane showing the nullclines for CycE and Rb intersecting at two stable (filled circles) and one unstable (unfilled circle) steady states when CycD is set to 0 $AU_D$. There is only one steady state at CycD = 0.15 $AU_D$ The system was reduced to two dimensions by making a steady state approximation for E2f and calculating E2f:Rb and pRb via conservation laws. **b** Bifurcation diagrams of the non-reduced system with constant E2F, showing stable (solid line) and unstable (dotted line) steady states of CycE. Line endings within the axes limits indicate disappearance of a steady state. **c** Time course of unphosphorylated Rb, CycE and E2f at CycD = 0.35 $AU_D$. For variable abbreviations please refer to Table 1. Parameter values and initial conditions are available the /versions/v0.0.1/ directory of the cell_cycle_model GitHub repository.

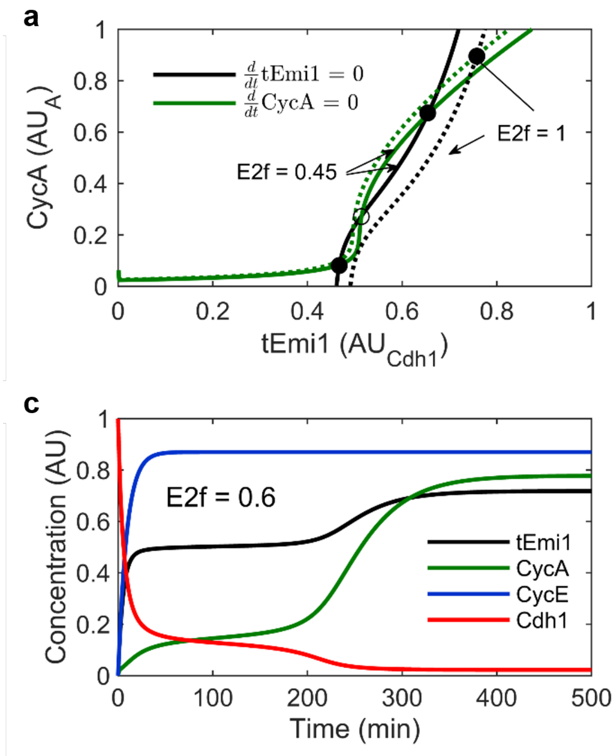


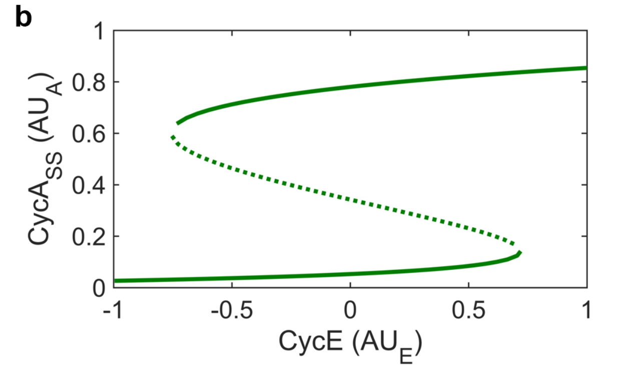


**Figure 2 — G1/S transition submodel.** **a** Phase plane showing the nullclines for tEmi1 and CycA intersecting at two stable (filled circles) and one unstable (unfilled circle) steady states when E2f is set to 0.45 $AU_{E2f}$. There is only one steady state at E2f = 1 $AU_{E2f}$. The system was reduced to the two plotted dimensions by making steady state approximations for the remaining variables. **b** Bifurcation diagram of the non-reduced system at E2f = 0.7 $AU_{E2f}$ with CycE as bifurcation parameter. **c** Time course of tEmi, CycA, CycE and Cdh1 at E2f = 0.6 $AU_{E2f}$. For the remaining variable abbreviations please refer to Table 1. Parameter values and initial conditions are available the /versions/v0.0.1/ directory of the cell_cycle_model GitHub repository.

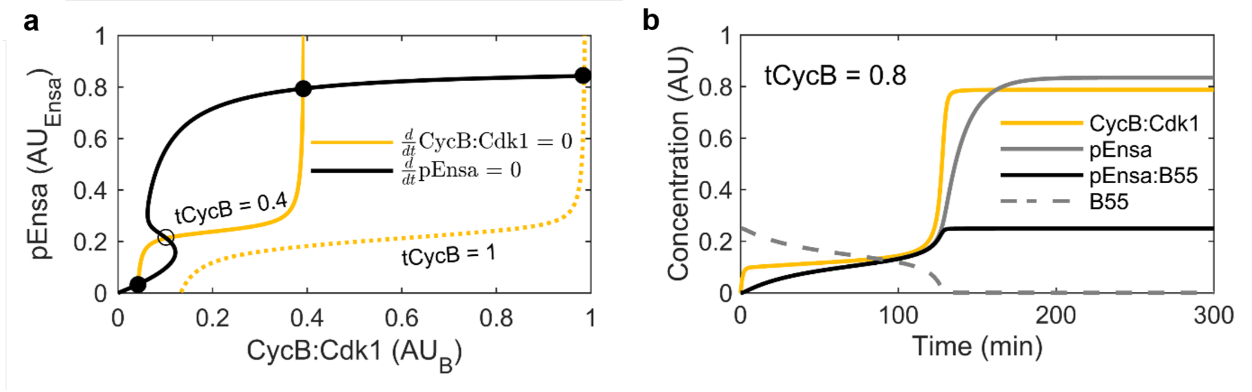


**Figure 3 — G2/M transition submodel as developed by Vinod and Novak [1]. a** Phase plane showing the nullclines for CycB:Cdk1 and pEnsa intersecting at two stable (filled circle) and one unstable (unfilled circle) steady states when tCycB is set to 0.4 $AU_B$. There is only one steady state at tCycB = 1 $AU_B$. The system was reduced to the two plotted dimensions by making steady state approximations for the remaining variables. Plot recreated from [1]. **b** Time course of CycB:Cdk1, pEnsa, pEnsa:B55, and B55 at tCycB = 0.8 $AU_B$. pWee1: phosphorylated Wee1; Cdc25: unphosphorylated Cdc25, Gw: unphosphorylated Greatwall kinase; Ensa: unphosphorylated Endosulfine alpha. For the remaining variable abbreviations please refer to Table 1. Parameter values and initial conditions are available from the /versions/v0.0.1/ directory of the cell_cycle_model GitHub repository.

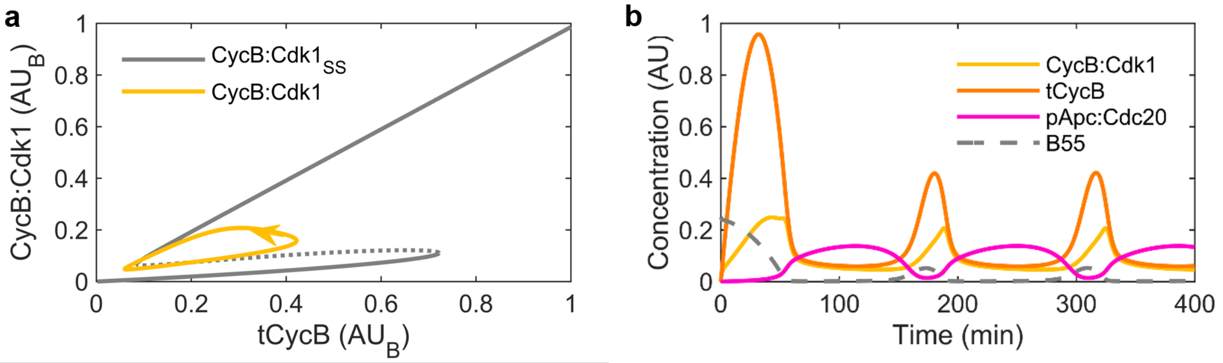


**Figure 4 — G2/M transition submodel with negative feedback of the M/A transition. a** CycB:Cdk1 steady states (grey) of the G2/M transition submodel shown in the bifurcation diagram of Figure 1f are overlaid with a stable limit cycle trajectory (yellow) of the system in the CycB:Cdk1 – tCycB subspace at a cyclin B synthesis rate of kSyCb $= 0.05\,\mathrm{min}^{-1}$. The periodic over- and undershooting of the upper and lower bifurcation point, respectively, results from combining the toggle switch of the G2/M submodel with the negative feedback of the M/A submodel. **b** Time course of CycB:Cdk1, tCycB, pApc:Cdc20 and B55 at kSyCb $= 0.05\,\mathrm{min}^{-1}$. For variable abbreviations please refer to Table 1. Parameter values and initial conditions are available the /versions/v0.0.1/ directory of the cell_cycle_model GitHub repository.

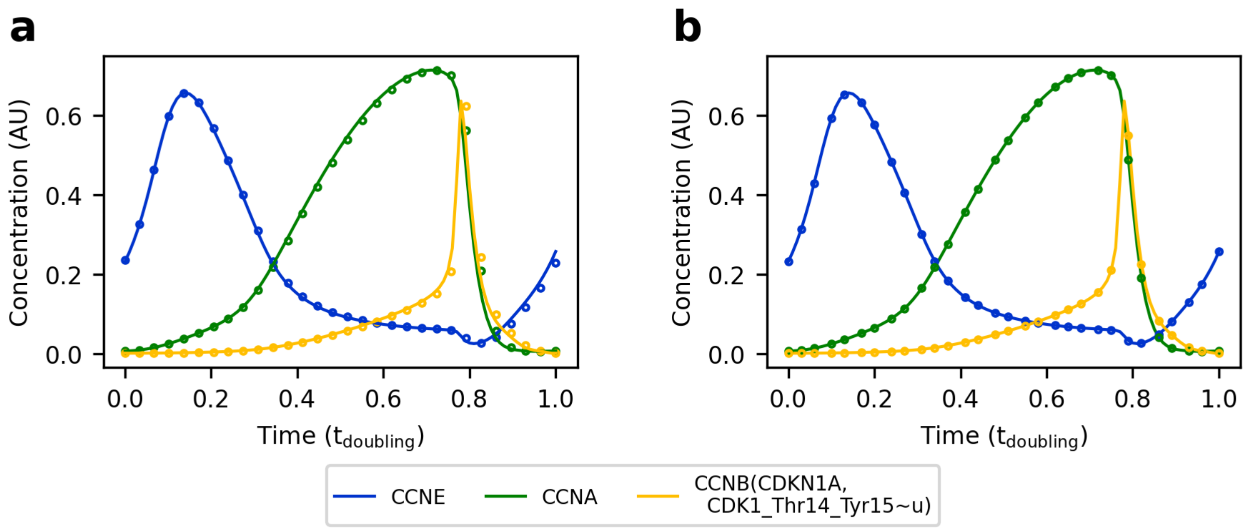


**Figure 5 — Comparison between model versions.** All simulations were performed with Copasi LSODA and default settings. Lines represent simulations of model version 2.1.4. **(a)** Circles represent second cycle of model version 1.0.0. First cycle is not shown, as the initial conditions of this model do not lie on the limit cycle trajectory. **(b)** Circles represent model version 3.0.0, simulated after export to SBML.

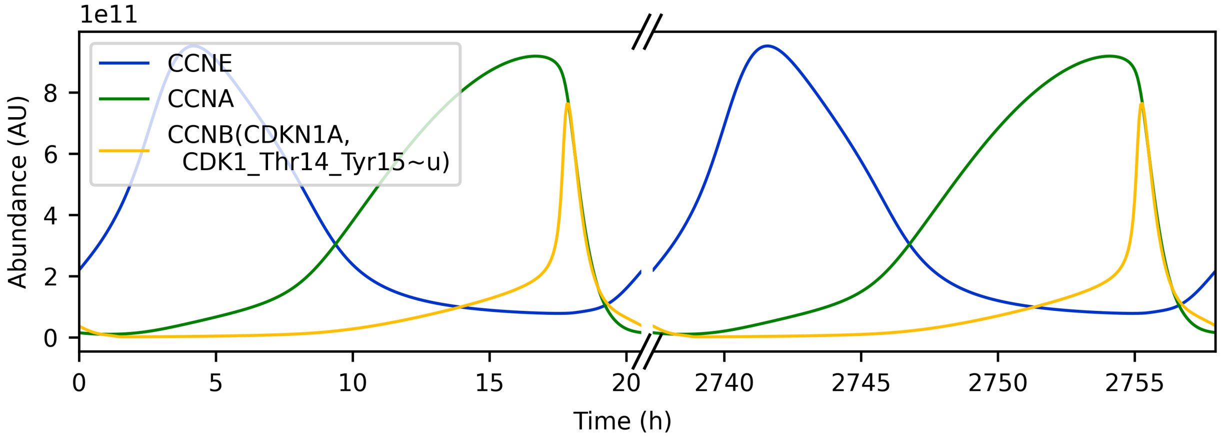


**Figure 6 — Cell cycle model with nuclear and cytoplasmic compartments.** Time course of three representative species over 2758 h. Model version 4.0.0.

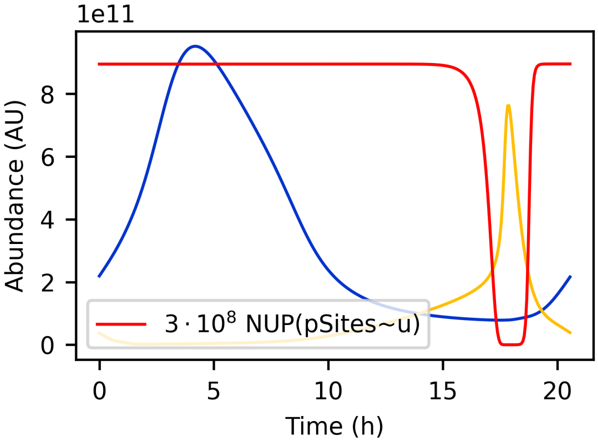


**Figure 7 — Cell cycle model with nuclear pore phosphorylation.** Time course of three representative species. Unphosphorylated nuclear pore complex `NUP(pSites~u)` are a proxy for an intact nuclear membrane. Model based on version 4.0.0 plus nuclear pore phosphorylation.

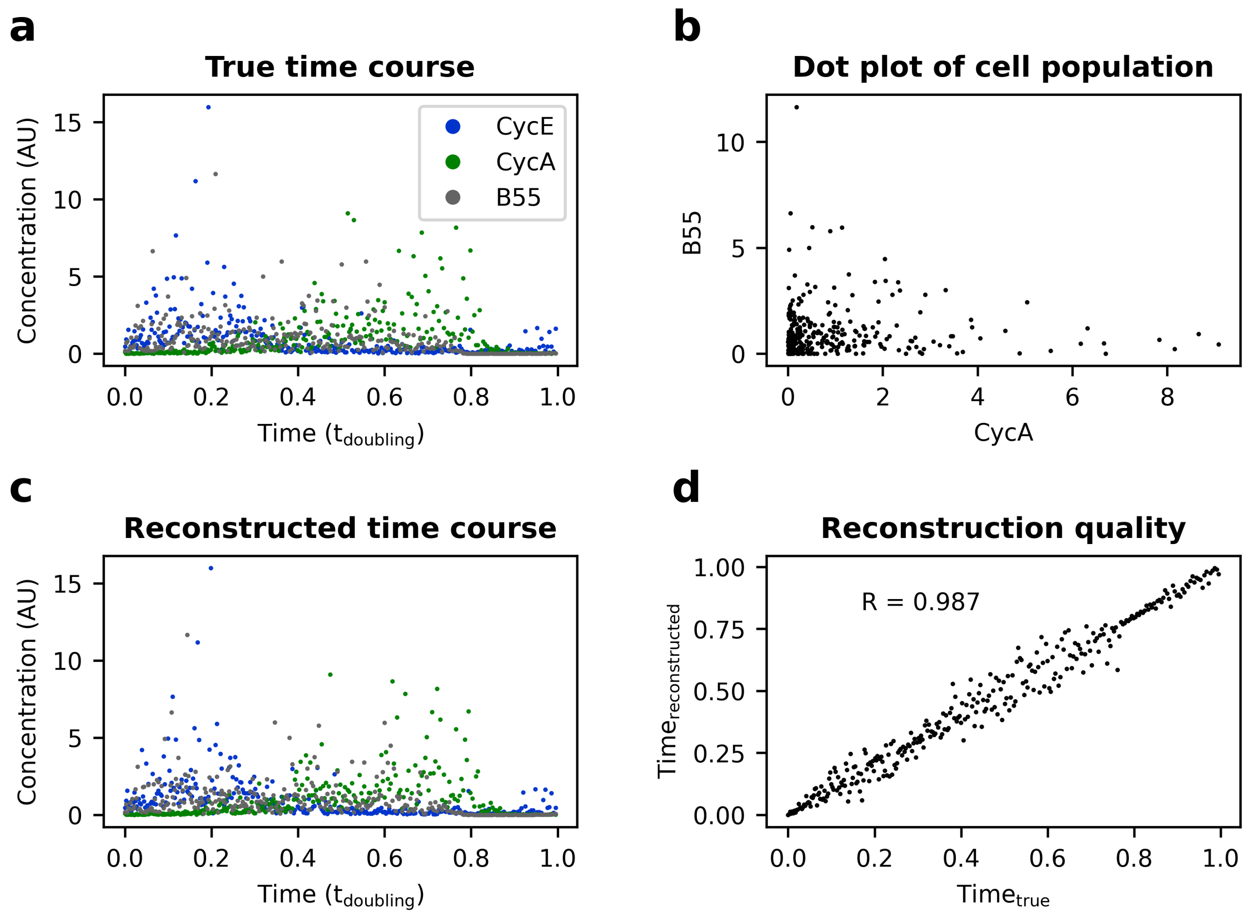


**Figure 8 — Cell cycle trajectory reconstruction from noise-free simulated data with reCAT. (a)** 300 cells were sampled across one cell cycle model simulation (version 2.1.4), such that cell density decreases exponentially over the cell cycle with one cell cycle length half-life. Of all 9 variables that were supplied to reCAT, only cyclin E, cyclin A and B55 are shown. **(b)**. Scatterplot of the sampled cells from A. **(c)** Reconstructed cell cycle trajectory. **(d)** Correlation between true and reconstructed cell cycle position for each cell.

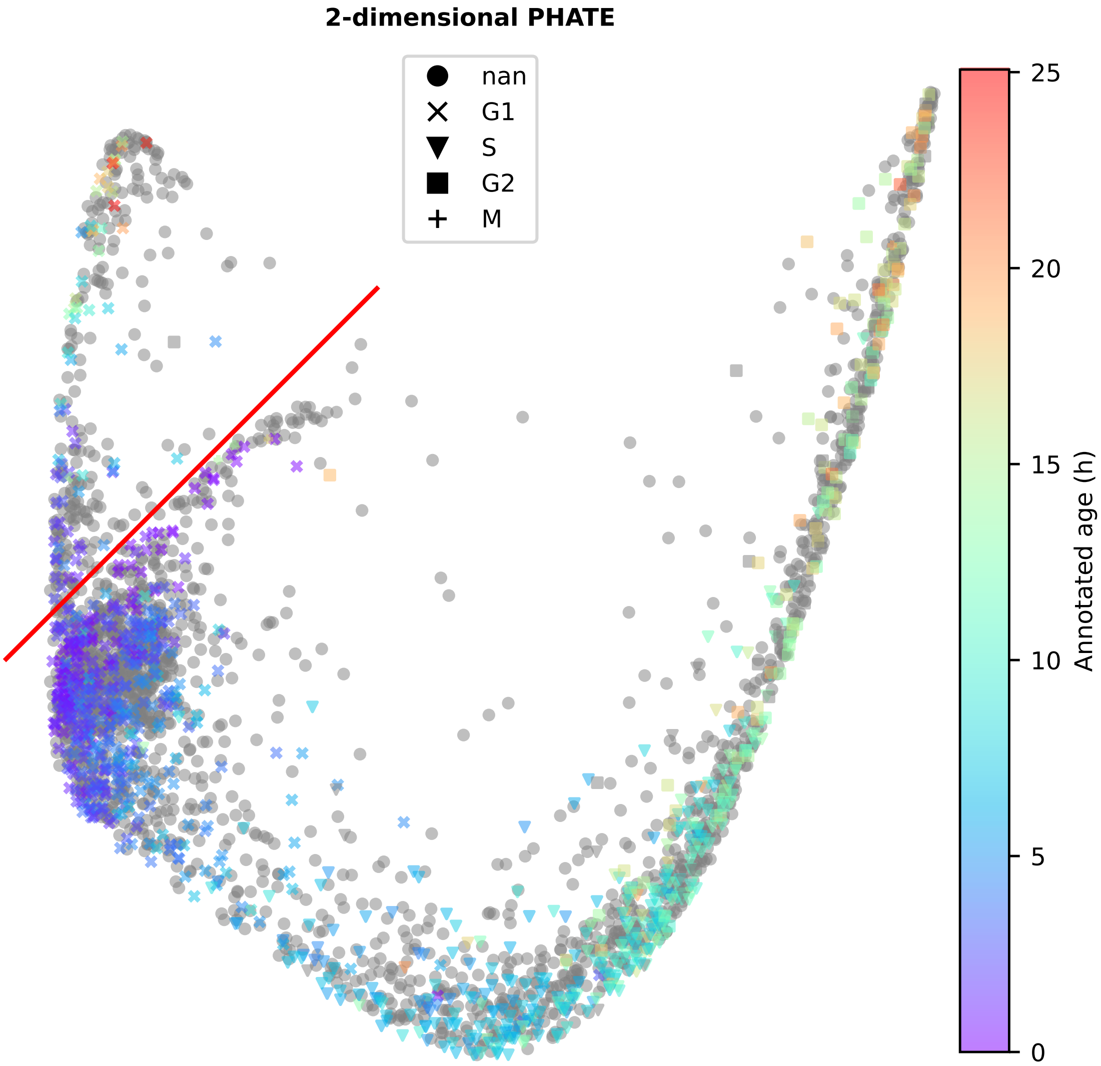


**Figure 9 — Gating proliferating cells using PHATE.** PHATE was performed on publicly available 4i measurements of an asynchronously dividing RPE-1 population [2]. Cells above the line indicated by the red line segment were classified as G0. For better visualisation only one third of all cells are shown.

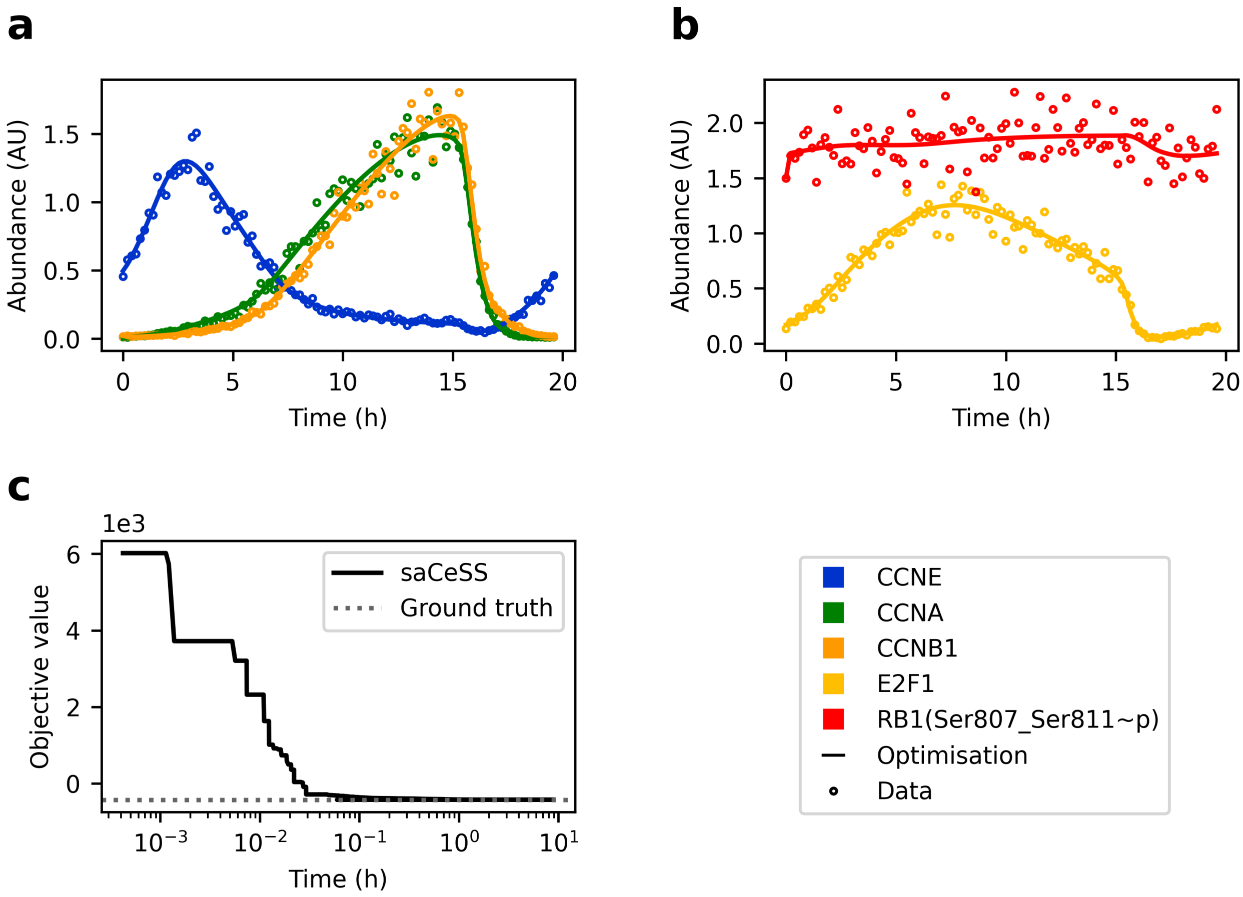


**Figure 10 — Testing self-adaptive Cooperative enhanced Scatter Search (saCeSS).** 300 datapoints of five observables with exponentially increasing spacing were generated from simulation results of model version 3.0.0. The datapoints were corrupted with $\mathcal{N}(1, 0.1^2)$ multiplicative noise (circles). saCeSS optimised the parameters from within a $[0.1 \cdot \boldsymbol{\theta}_{true}, 10 \cdot \boldsymbol{\theta}_{true}]$. **a, b** The lines show simulated time courses for the observables, using the parameter set found by saCeSS (legend in lower right panel). **c** Convergence curve of the negative log-likelihood during optimisation using additive measurement noise ($\sigma = 0.1$). Initial parameter guess was ignored).

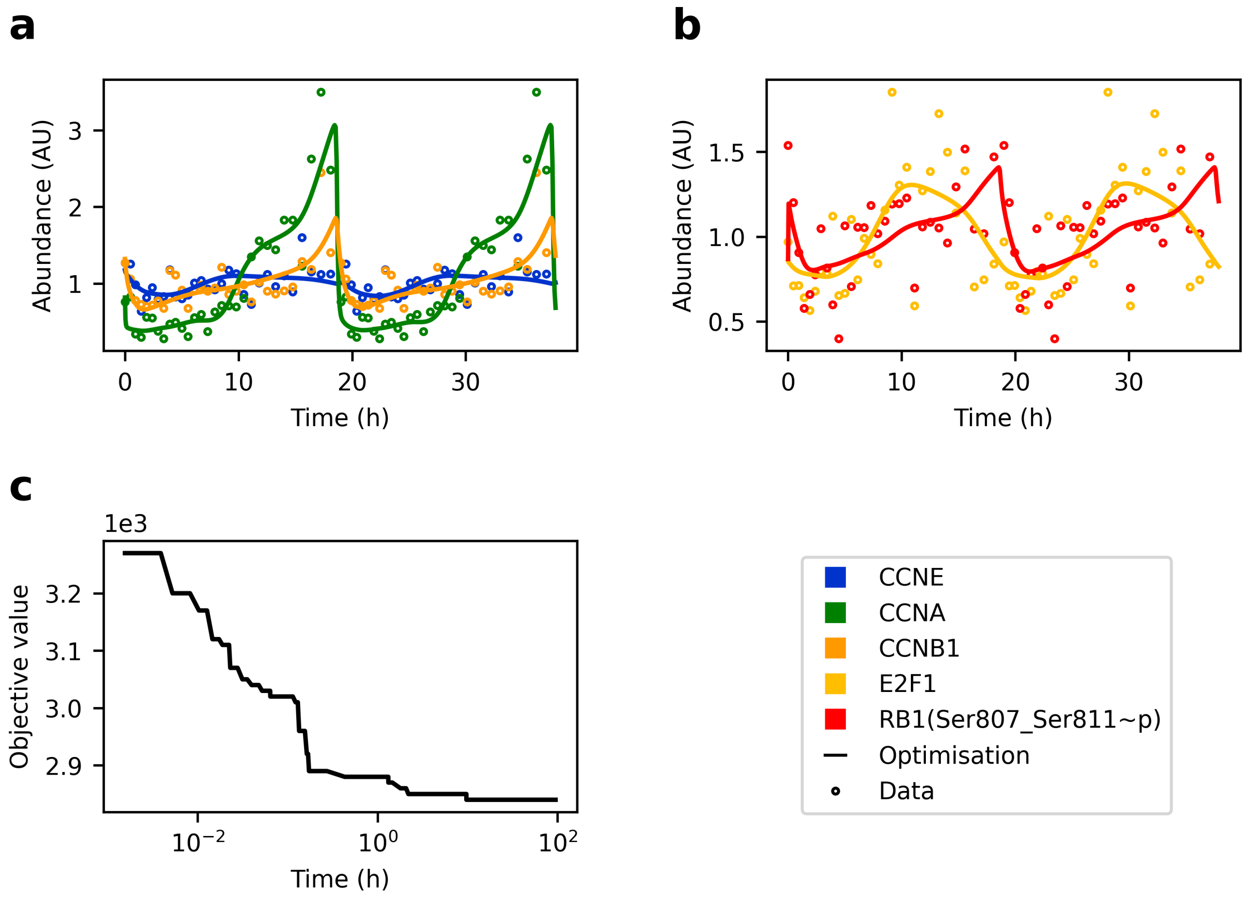


**Figure 11 — Model version 3.0.1 fitted to pseudo-time courses of RPE-1 cells. (A, B)** Experimental measurements and simulation results using estimated parameters. For better visibility, only every $10^{\text{th}}$ measurement is shown. **(C)** Convergence curve of the negative log-posterior during optimisation using additive measurement noise ($\sigma = 1$). Initial parameter guess was provided (see PEtab parameter table).

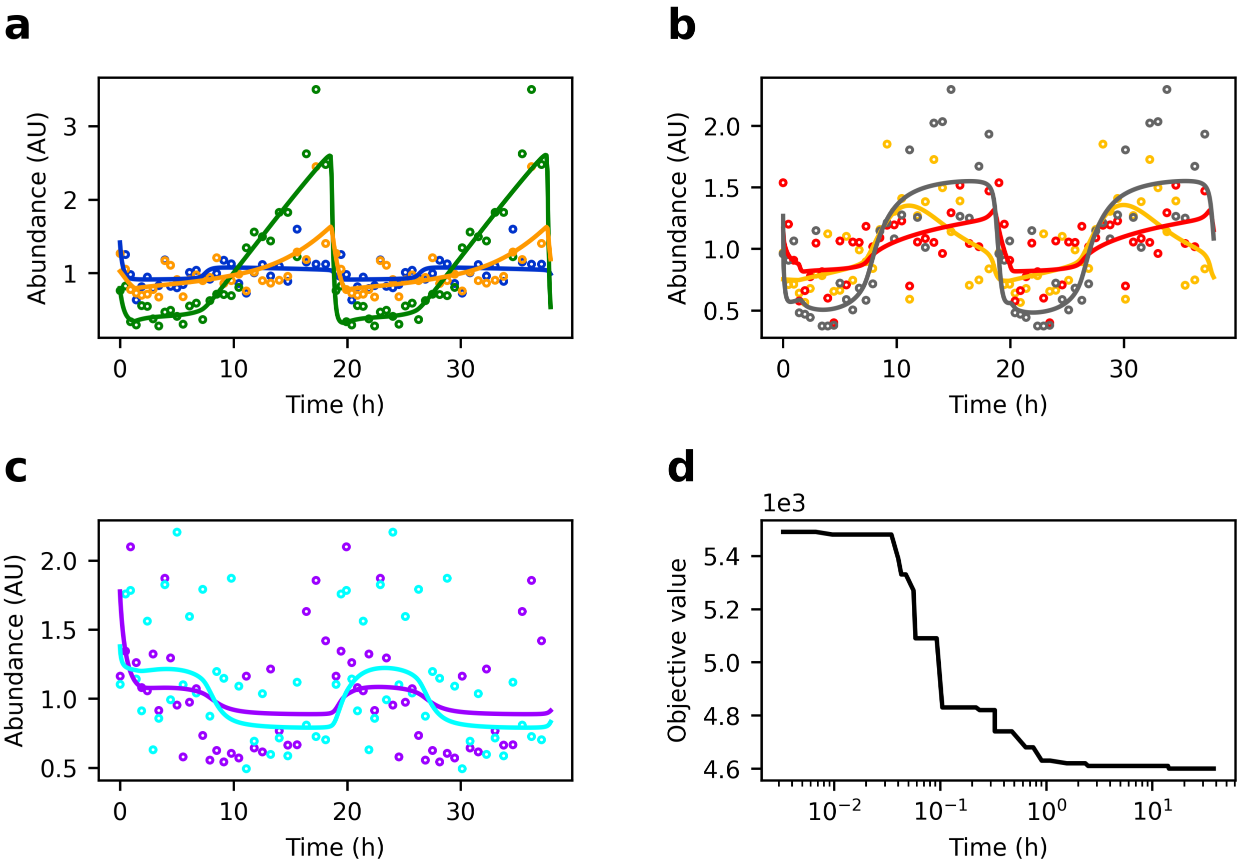


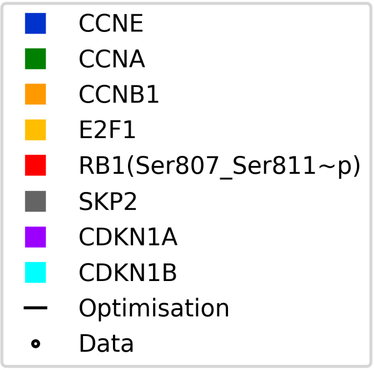


**Figure 12 — Model version 3.2.0 fitted to pseudo-time courses of RPE-1 cells.** **(a–c)** Experimental measurements and simulation results using estimated parameters. For better visibility, only every $10^{\text{th}}$ measurement is shown. **(d)** Convergence curve of the negative log-posterior during optimisation using additive measurement noise ($\sigma = 1$). Initial parameter guess was provided to the optimiser (see PEtab parameter table).

### Supplementary Note 1: Merging submodels

When merging the submodels to a full cell cycle model, we had to introduce (a) new reactions (e.g.: CycA mediated Rb phosphorylation; Apc:Cdh1 mediated CycB degradation) and (b) new species (e.g.: pApc:Cdh). To connect the G1/S submodel with the G2/M submodel, we also introduced the transcription factor FOXM1. FOXM1 is synthesized by E2f and activated by Cdks [3]. It binds to the promoter of cyclin B to promote its synthesis [4]. Furthermore, certain molecules that had accumulated during earlier cell cycle stages (tE2F, tEmi1, CycA, tFoxM1 tCdc20) still had to be degraded. For the SCF substrates tE2f [5] and tEmi1 [6] (but for simplicity not for CycE and CycA) this was accomplished by introducing a new, phosphorylated species that was targeted for ubiquitination. These species are phosphorylated by CycA and CycB, thus introducing additional negative feedback loops on E2F (e.g.: E2F activates CycA, which inhibits E2F; and E2F activates CycB via CycA mediated FOXM1 phosphorylation, and CycB inhibits E2F) and on Emi1 (e.g.: Emi1 inhibits Apc:Cdh1 inhibits CycA/B inhibits Emi1). For the (p)Apc:Cdh1 substrates tFoxM1 [7] and tCdc20 [8] a (p)Apc:Cdh1 mediated degradation reaction was introduced. This again results in new feedback loops. For instance, FoxM1 activates CycB. CycB and (p)Apc:Cdh1 mutually inhibit each other. (p)Apc:Cdh1 then inhibits FoxM1. The (p)Apc:Cdh1 mediated degradation of tFoxM1 ensures that tCycB is kept low in G1 phase. Similarly, for CycA/B a pApc:Cdc20 mediated degradation reaction was introduced, resulting in yet another negative feedback loop (CycA inhibits (p)Apc:Cdh1, inhibits pApc:Cdc20, inhibits CycA). These changes also required modifications in the parameters to preserve characteristic features of each submodel:

- RP: The restriction point is a point after which cell cycle progression is mitogen independent.
- G1/S: There is a delay between CycE and CycA accumulation.
- G2/M: There is a sharp increase in CycB:Cdk1.
- M/A: Cyclin B is rapidly and almost completely degraded.

To reduce the complexity of this task, the submodels were fused in a stepwise manner. The resulting model of the whole cell cycle (cell_cycle_v1.0.0) comprised 37 species (Table 1). The corresponding equations are available in file /versions/v1.0.0/cell_cycle_v1.0.0.ode of the cell_cycle_model GitHub repository.

### Supplementary Note 2: Naming conventions in the BioNetGen model

The BNGL syntax also lends itself well to add semantic meaning to the model variables. This enables to conceive the following naming convention:

- Proteins
  - Protein molecule types were named by there Human Genome Organization Gene Nomenclature Committee (HGNC) short name. No distinction is made between protein isoforms that are transcribed from the same gene. Example: `RB1()`.
  - Protein family types were described by a shortened protein HGNC name where unambiguously possible. Example: the set of CCNE1 and CCNE2 was collectively referred to as `CCNE()`. Otherwise, they were described as the concatenation of HGNC names separated by underscores. Example: `ENSA_ARPP19()`.
- Promoters
  - Promoters where described by the protein they activate, suffixed by `_promoter`. Example: `CDC20_promoter()`.
- Binding sites
  - If the binding site is bound by a protein or protein family, the binding site is named by the protein (family) name. Example: `PPP2R2B(ENSA_ARPP19)`.
  - If the binding site is bound by a promoter, the binding site is named `DBD` (short for DNA binding domain). Example: `FOXM1(DBD)`.
- Phosphorylation sites
  - The unphosphorylated state is indicated with a `u`, the phosphorylated state with a `p`.
  - Single site phosphorylation were named by the three letter amino acid code, suffixed with the amino acid position. Example: `MASTL(Thr198~u~p)`.
  - Known multisite phosphorylations were named by the concatenation of the single site phosphorylation names, separated with an underscore. Example `RB1(Ser807_Ser811~u~p)`. Note that no comma separates `Ser807` and `Ser811` in this example. Therefore, `Ser807_Ser811` is treated as a single site.
  - Unknown phosphorylation sites could have any name.
    Example `FZR1(APC,FBXO5,nTerm~u~p)`.
- Exceptions
  - `E2F()` means only the activating E2F transcription factors E2F1, E2F2 and E2F3.
  - The anaphase-promoting complex was named `APC()`.
  - The nuclear pore complex was named `NUP()`.

The exceptions indicate that some generalisability was traded off against concise syntax. Of note, as much as any biochemical model is incomplete, the use of the above syntax shall not imply that no other phosphorylation sites, binding sites or molecule types than those explicitly described by the model are involved in the real phenomenon of cell cycle control. Rather, it shall just help to describe more precisely which biological phenomena are considered in the model and which are not.

### Supplementary Note 3: Adding CDKN1B to the cell cycle model

Like CDKN1A, the CDKN1B cycling kinase inhibitor, also known as p27, confers ultrasensitivity. Together, the RP, the G1/S transition, the DNA damage checkpoint, and the mutual inhibition between CDKN1B and cyclin:Cdk complexes represent four partially redundant mechanisms to ensure bistability in G1. Redundancy confers robustness in varying or perturbed conditions. As robustness under perturbation seems to be an important feature of the cell cycle, CDKN1B was added to the model, along with FOXO transcription factors which activate its expression [9]. Unlike CDKN1A, CDKN1B does not bind cyclin B:Cdk complexes [10]. There is also clear evidence that Thr187 phosphorylation not only prevents cyclin:Cdk binding, but also targets CDKN1B for SKP2 mediated degradation.

### Supplementary Note 4

To implement nucleocytoplasmic shuttling, the following points were considered:

- The nucleocytoplasmic transport reactions described in the literature may only represent a subset of the nucleocytoplasmic transport reactions that are happening in the cell.
- We aim at identifying an extensive biochemical reaction network for which a parameterisation exists, such that the network is capable of explaining the regulation of the full cell cycle.
- We aim to demonstrate that the model can generate sustained oscillations with manually chosen parameters.

In agreement with these considerations, translocation reactions for all molecular species, except for those that are exclusively located in either compartment (i.e. promoters), were added to the model. The kinetic constants for the transport reactions were set to very large and equal values (to be later constrained during parameter estimation). This ensured that the oscillatory behaviour of single-compartment versions of the model was retained. More specifically, the following adaptations were made to version `3.2.0`:

- Four compartments were defined using BNGL: a $1 \times 10^{-7}$ dm thick plasma membrane (`Plm`, $2 \times 10^{-14}$ dm$^3$), cytoplasm (`Cyt`, $1 \times 10^{-12}$ dm$^3$), a $1 \times 10^{-7}$ dm thick nuclear membrane (`Num`, $9 \times 10^{-15}$ dm$^3$) and a nucleus (`Nuc`, $1 \times 10^{-12}$ dm$^3$).
- Nuclear pores `NUP(pSites~u~p)` were inserted into the nuclear membrane.
- Instead of describing the complete Ran-GTP pathway, simple import and export reactions through unphosphorylated nuclear pores were added for all species that do not contain promoters. The kinetic constants for import and export were set to same very large value to allow quasi-free diffusion of molecules.
- The initial concentration of promoters was set to zero in the cytoplasm and a positive value in the nucleus.
- Reaction rules were made independent of localisation. I.e. reactions are unaware of the compartment they occur in, use the same kinetic constants independent of the compartment, and fire at any location where all reactants are present. Only protein synthesis reactions contain a notion of localisation, namely that proteins are synthesized into the cytoplasm.
- Nuclear envelope breakdown was imitated by introducing a rule for `CCNB(CDKN1A,CDK1_Thr14_Tyr15~u)` mediated nuclear pore phosphorylation, allowing for quasi-free diffusion of all pure protein species.

The above changes lead to model version `4.0.0`. Simulation results are shown in Supplementary Fig. 7. However, due to stiffness arising from nuclear envelope breakdown, we switched it off throughout the present study.

### Supplementary Note 5

Stallaert *et al.* [2] found that a substantial fraction of RPE-1 cells exit the proliferative cell cycle trajectory into a non-proliferative G0 arm. Removing such cells results in a purely proliferative trajectory. However, it is unclear whether the distribution of the cells along the trajectory follows the pseudo-time expressed by Equation 1 in the main text. This is because G0 cells may not enter the proliferative cell cycle at the same point of the trajectory where they left. Should they do parts of the proliferative trajectory twice, the calculated pseudo-time would become artificially stretched in these regions. Conversely, should they skip parts of the proliferative trajectory, the calculated pseudo-time would become artificially compressed. It is clear that repeating mitosis and skipping parts of S-phase via a G0 trajectory cannot be the norm, as this would lead to regular chromosome aberrations. Nevertheless, former G0 cells may move along a trajectory that is mostly parallel to the proliferative trajectory, but shifted in a few states for some time, before completely merging with the proliferative trajectory. Such parallel trajectories would bias the reconstructed trajectory in the direction of the shift. Yet, overall the trajectory reconstruction would only be strongly affected if alternative trajectories skip or redo large stretches of the proliferative trajectory and a large amount of cells at any given time would move on these trajectories. As both can be considered highly speculative at present, it seems reasonable to assume that removal of G0 cells from an asynchronous cell population leads to a quasi asynchronous cell population that mostly moves along the proliferative cell cycle trajectory. That is, the cell population can be used for reconstructing a proliferative trajectory from snapshot data and for calculating pseudo-time from ranks via Equation 1.

### Supplementary Note 6

Using real experimental data instead of an artificial dataset required us to make five minor modifications to the optimisation strategy. First, simulations with the optimised parameters did not result in oscillations. By duplicating the experimental data to obtain two consecutive cycles, oscillatory behaviour could be enforced. Second, simulations with these optimised parameters unexpectedly showed small dampened oscillations of observables for a short period of time, just after the metaphase anaphase transition. Attempts of manually adjusting the parameters led to the following hypothesis. Residual CCNA and CCNB1 fluorescence signal after anaphase in the Stallaert dataset enforce too high levels of these two species in the model. As a consequence, there is substantial E2F1 phosphorylation, and thus degradation in the model, especially when the activity of the counteracting phosphatase PPP2R2B is suppressed by action of CCNB1. To overcome these high degradation levels, E2F1 synthesis rates must be high. In other words, E2F abundance is highly responsive to changes in kinase/phosphatase ratio. Once kinase/phosphatase ratio drops, E2F1 shoots up, activates slow CCNA synthesis, which results in E2F1 phosphorylation and thus degradation with a delay. Attempts to manually reduce E2F1 degradation and synthesis rates failed to produce sustained oscillations. However, the active concentration of cyclins may be lower than the antibody signal suggests, for instance due to incomplete background subtraction in the dataset. To account this possibility, the offset parameters $o_k$ in the observation model defined in Equation (2) of the main text were allowed to deviate from zero. Third, using real world data, the global optimum may lie out of the specified parameter bounds. Therefore, active windows (i.e. windows where the best possible solution found by the optimiser lies at a boundary) were shifted in the corresponding direction by a factor of 10. These shifts were performed iteratively between the multiple job submissions to that were required due to limitations of maximal job execution times on the cluster. Fourth, saCeSS tended to perform better with DHC than parPE/ipopt as local solver option. Last, to further incorporate knowledge that cyclins are present in higher concentrations than E2F1 and RB1, Laplacian priors were introduced for the scaling parameters of the cyclins.

### References


[1] P. K. Vinod and Bela Novak. "Model scenarios for switch-like mitotic transitions". *FEBS Letters* 589.6 (2015), pp. 667–671.

[2] Wayne Stallaert et al. "The structure of the human cell cycle". *Cell Systems* 0.0 (2021).

[3] J. Millour et al. "ATM and p53 regulate FOXM1 expression via E2F in breast cancer epirubicin treatment and resistance". *Mol Cancer Ther* 10.6 (2011), pp. 1046–58.

[4] H. Murakami et al. "Regulation of yeast forkhead transcription factors and FoxM1 by cyclin-dependent and polo-like kinases". *Cell Cycle* 9.16 (2010), pp. 3233–42.

[5] A. Marti et al. "Interaction between ubiquitin–protein ligase SCFand E2F-1 underlies the regulation of E2F-1 degradation". *Nature Cell Biology* 1.1 (1999), p. 14.

[6] Daniele Guardavaccaro et al. "Control of Meiotic and Mitotic Progression by the F Box Protein $\beta$-Trcp1 In Vivo". *Developmental Cell* 4.6 (2003), pp. 799–812.

[7] J. Laoukili et al. "FoxM1 is degraded at mitotic exit in a Cdh1-dependent manner". *Cell Cycle* 7.17 (2008), pp. 2720–6.

[8] C. M. Pfleger and M. W. Kirschner. "The KEN box: an APC recognition signal distinct from the D box targeted by Cdh1". *Genes Dev* 14.6 (2000), pp. 655–65.

[9] Fangfang Yang et al. "The Akt/FoxO/p27Kip1 axis contributes to the anti-proliferation of pentoxifylline in hypertrophic scars". *Journal of Cellular and Molecular Medicine* 23.9 (2019), pp. 6164–6172.

[10] Alessia Montagnoli et al. "Ubiquitination of p27 is regulated by Cdk-dependent phosphorylation and trimeric complex formation". *Genes & Development* 13.9 (1999), pp. 1181–1189.
